# Enzymatic reactions of AGO4 in RNA-directed DNA methylation: siRNA duplex loading, passenger strand elimination, target RNA slicing and sliced target retention

**DOI:** 10.1101/2022.10.06.511223

**Authors:** Feng Wang, Hsiao-Yun Huang, Jie Huang, Jasleen Singh, Craig S. Pikaard

**Affiliations:** Department of Biology and Department of Molecular and Cellular Biochemistry, Indiana University, Bloomington, IN 47405, USA; Howard Hughes Medical Institute, Indiana University, Bloomington, IN 47405, USA

**Keywords:** ARGONAUTE, short-interfering RNA, noncoding RNA, transcriptional gene silencing, chromatin modification, RNA Polymerase V

## Abstract

RNA-directed DNA methylation in plants is guided by 24 nt siRNAs generated in parallel with 23 nt RNAs of unknown function. We show that 23 nt RNAs function as passenger strands during 24 nt siRNA incorporation into AGO4. The 23 nt RNAs are then sliced into 11 and 12 nt fragments, with 12 nt fragments remaining associated with AGO4. Slicing recapitulated with recombinant AGO4 and synthetic RNAs reveals that siRNAs of 21-24 nt, with any 5’ terminal nucleotide, can guide slicing, with sliced RNAs then retained by AGO4. In vivo, RdDM target locus RNAs that copurify with AGO4 also display a sequence signature of slicing. Comparing plants expressing slicing-competent versus slicing-defective AGO4 shows that slicing elevates cytosine methylation levels at virtually all RdDM loci. We propose that siRNA passenger strand elimination and AGO4 tethering to sliced target RNAs are distinct modes by which AGO4 slicing enhances RNA-directed DNA methylation.

## Introduction

In eukaryotes, RNA silencing plays a significant role in gene regulation, transposable element repression and genome stability. A common theme among RNA silencing pathways is that small RNAs stably associate with Argonaute family proteins and then interact with complementary target RNAs to interfere with RNA translation or to bring about chromatin modifications that repress gene transcription (Martienssen and Moazed, 2015; Iwakawa and Tomari, 2022). An ancestral and fundamental function shared by many Argonaute proteins is small RNA-programmed endonucleolytic slicing of the targeted RNA at a phosphodiester bond (Joshua-Tor and Hannon 2011).

RNA-directed DNA methylation (RdDM) is the major transcriptional gene silencing pathway in plants (Matzke and Mosher 2014; Zhou and Law 2015; Wendte and Pikaard 2017; Zhang et al. 2018). RdDM siRNA precursors are synthesized by DNA-dependent NUCLEAR RNA POLYMERASE IV (Pol IV) (Ream et al. 2009; Haag and Pikaard 2011) and RNA-DEPENDENT RNA POLYMERASE 2 (RDR2) (Zhou and Law 2015; Fukudome et al. 2021; Du et al. 2022). These enzymes physically associate and carry out coupled reactions that transcribe target locus DNA sequences into double-stranded RNAs of ∼25-40bp (Haag et al. 2012; Singh et al. 2019; Fukudome et al. 2021; Huang et al. 2021). DICER-LIKE 3 (DCL3) then cuts the dsRNAs into siRNAs, guided by sequence and structural features at the ends of precursor duplexes (Loffer et al. 2022). Resulting dicing products consist of a 24 nt siRNA strand paired with either a 23 nt strand (24/23 duplexes) or a 24 nt strand (24/24 duplexes) (Nagano et al. 2014; Loffer et al. 2022), one strand of which ultimately becomes stably associated with ARGONAUTE 4 (AGO4). The siRNAs then guide AGO4 to RdDM target loci by basepairing to transcripts generated by NUCLEAR RNA POLYMERASE V (Pol V) (Wierzbicki et al. 2008; Wierzbicki et al. 2009; Liu et al. 2018), with protein-protein interactions between AGO4 and the Pol V largest subunit (NRPE1), and/or the Pol V-associated protein, SPT5L also implicated in target locus interactions (Li et al. 2006; El-Shami et al. 2007; Bies-Etheve et al. 2009). Resulting AGO4-siRNA-Pol V complexes enable the recruitment of the *de novo* cytosine methyltransferase, DRM2 (Zhong et al. 2014; Fang et al. 2021) as well as enzymes that reposition or chemically modify the histone proteins that wrap the DNA (Du et al. 2015). Collectively, these activities bring about chromatin environments that repress promoter-dependent gene transcription by RNA Polymerases I, II, or III (Preuss et al. 2008; Blevins et al. 2009) (Figure 1A).

**Figure 1.**
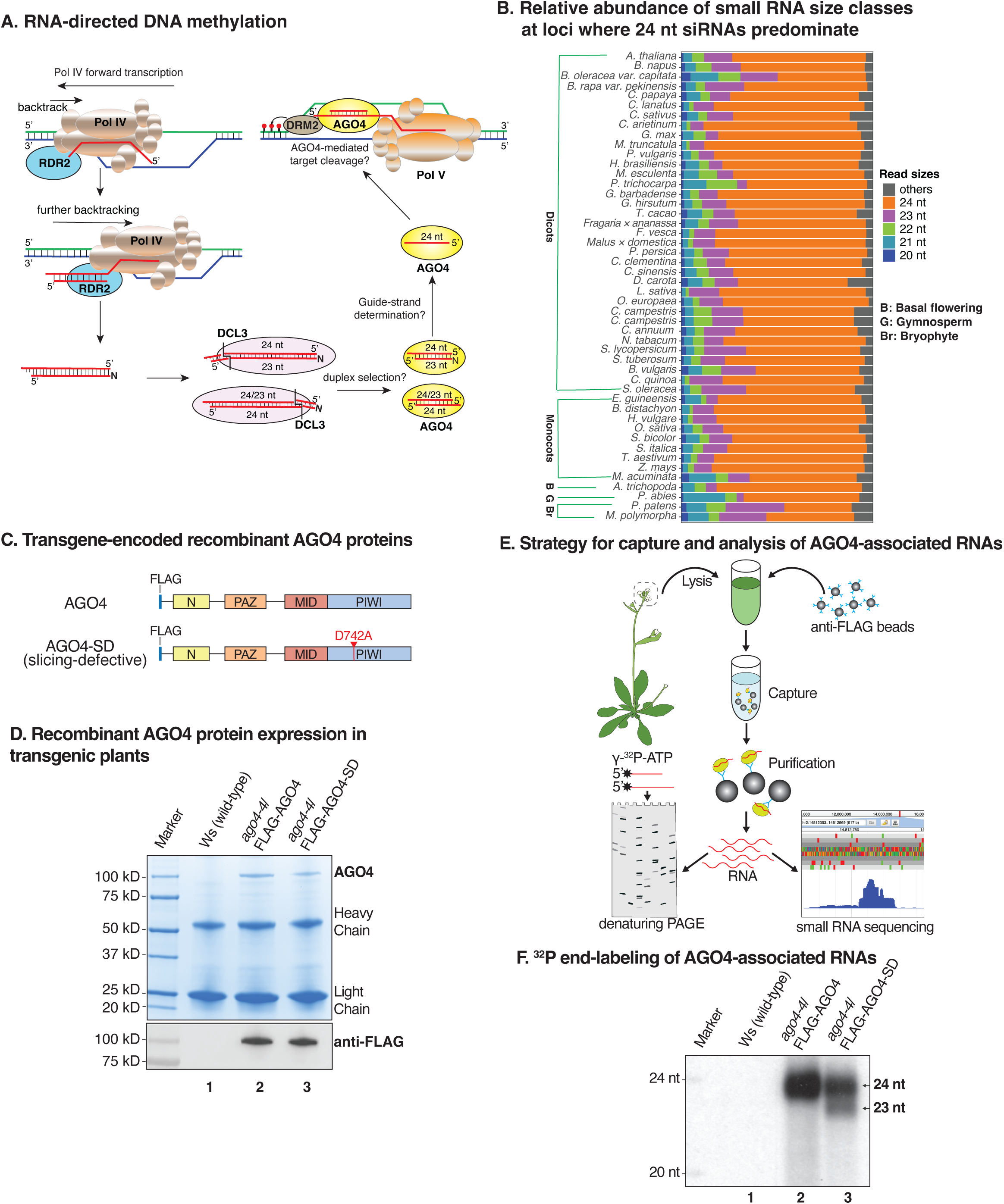
Evidence that 23 nt siRNAs function as passenger strands for 24 nt guide RNAs. **A.** A simplified model for RNA-directed DNA methylation highlighting questions addressed in our study. **B**. Relative abundance of small RNA size classes at 24 nt siRNA-dominated loci in diverse species (Lunardon et al. 2020). **C**. Transgene-encoded AGO4 proteins expressed from the native promoter in *ago4-4* null mutant plants (ecotype Ws). **D**. Comparison of AGO4 and AGO4-SD levels IPed from inflorescence tissue. Duplicate gels were stained with Coomassie Brilliant Blue (top panel) or subjected to immunoblotting using anti-FLAG antibody (bottom panel). Non-transgenic Ws serves as a control in lane 1. **E**. Cartoon depicting AGO4 affinity capture and analysis of associated RNAs by ^32^P end-labeling or deep sequencing. **F**. Autoradiogram of 5’ end-labeled RNAs in anti-FLAG IP fractions from non-transgenic plants (lane 1), plants expressing FLAG-AGO4 (lane 2) or plants expressing FLAG-AGO4-SD (lane 3).

Although DCL3 dicing frequently generates duplexes consisting of 23 nt siRNAs basepaired with 24 nt siRNAs (Loffer et al. 2022), siRNAs that co-immunoprecipitate with AGO4 are almost exclusively 24 nt (Qi et al. 2006; Havecker et al. 2010; Wang et al. 2011). Thus, the fates and functions of 23 nt siRNAs is unclear. Using genetic and genomic approaches, and enzymatic reactions catalyzed by recombinant AGO4-siRNA complexes reconstituted *in vitro*, we show that 23 nt siRNAs serve as passenger strands that specify that the 24 nt RNAs to which they are basepaired become stably associated with AGO4 as guide strands. The 23 nt RNAs are then sliced by AGO4 into 11 and 12 nt fragments. We show that AGO4 loading and slicing activity is similar for guide RNAs with A, U, C or G at their 5’ termini and for guide RNAs that vary in length from 21-24 nt, indicating that the 24 nt length and 5’A bias among AGO4-associated siRNAs *in vivo* is not due to AGO4 binding requirements. Surprisingly, we find that AGO4 retains fragments of the passenger strand and target RNAs that it slices, both *in vitro* and *in vivo*, suggesting a function for the retained RNAs. We propose that AGO4 slicing of target RNAs synthesized by Pol V elongation complexes causes the successive release of RNA fragments that remain locally associated with RdDM loci. By remaining bound to these sliced RNAs, multiple independent AGO4-RNA complexes could linger at the loci and act in parallel to achieve maximal levels of methylation.

## Results

### Plant 23 nt RNAs function as passenger strands for 24 nt guide RNAs

DCL3 dicing of double-stranded Pol IV-RDR2 transcripts yields duplexes with two 24 nt siRNAs (24/24 duplexes) or a 24 nt siRNA paired with a 23 nt siRNA (24/23 duplexes) (Loffer et al. 2022)(Figure 1A). Thus, at loci where 24 nt siRNAs are the predominant class of small RNA, in diverse plant species, 23 nt siRNAs are second in abundance (Figure 1B). Small RNAs of 21 and 22 nt are also produced at these loci, ostensibly diced by DCL 1, 2 or 4, and could potentially facilitate post-transcriptional silencing and/or initial low-level DNA methylation events that enable Pol IV and Pol V recruitment, thus jump-starting the major RdDM pathway diagrammed in Figure 1A (McCue et al. 2015; Hung and Slotkin 2021; Sigman et al. 2021).

Insight into the function of 23 nt siRNAs came from an experiment in which we transformed an *ago4-4* null mutant of the *A. thaliana* ecotype (natural strain) Wassilewskija (Ws) with transgenes expressing wild-type AGO4 or slicing-defective AGO4 (AGO4-SD), each bearing a FLAG epitope tag (Figure 1C). In AGO4-SD, aspartate D742 is changed to alanine within the catalytic center conserved among slicing-competent Argonaute family proteins (Rivas et al. 2005). The recombinant AGO4 and AGO4-SD proteins were expressed at similar levels, as shown by anti-FLAG immunoblot detection of the proteins in total cell lysates (Figure S1A), and Coomassie-blue staining and anti-FLAG immunoblot detection following their affinity capture using anti-FLAG antibodies (Figure 1D). RNAs co-purifying with affinity-captured AGO4 or AGO4-SD were then 5’ end-labeled with ^32^P and resolved by denaturing polyacrylamide gel electrophoresis (PAGE) or subjected to high-throughput small RNA sequencing (Figure 1E).

End-labeling showed that small RNAs associated with wild-type AGO4 are almost exclusively 24 nt (Figure 1F, lane 2), consistent with prior studies (Qi et al. 2006; Havecker et al. 2010; Wang et al. 2011). However, siRNAs associated with slicing-defective AGO4-SD included both 24 nt and 23 nt RNAs (Figure 1F, lane 3) at relative levels consistent with their biogenesis (Singh et al. 2019; Loffer et al. 2022). These results suggest that AGO4 is initially loaded with siRNA duplexes, with one strand (the passenger strand) then eliminated by slicing (Ye et al. 2012). For 24/23 duplexes, the 23 nt strand is selectively eliminated.

### AGO4-associated small RNAs have distinct sequence features that allow interpretation of their origins

Deep sequencing of small RNAs that copurify with FLAG-tagged wild-type AGO4 upon anti-FLAG immunoprecipitation (IP) revealed abundant 24 nt RNAs and minor classes of 12, 23 and 25 nt RNAs (Figure 2A; Figure S1B). By contrast, RNAs associated with slicing-defective AGO4-SD included abundant 24 and 23 nt RNAs (Figure 2A, Figure S1B) and low levels of 25 and 26 nt RNAs, but no 12 nt RNAs. Non-transgenic plant samples subjected to anti-FLAG immunoprecipitation as a negative control yielded no RNA signals above background (Figure 2A, bottom panel labeled Mock IP).

**Figure 2.**
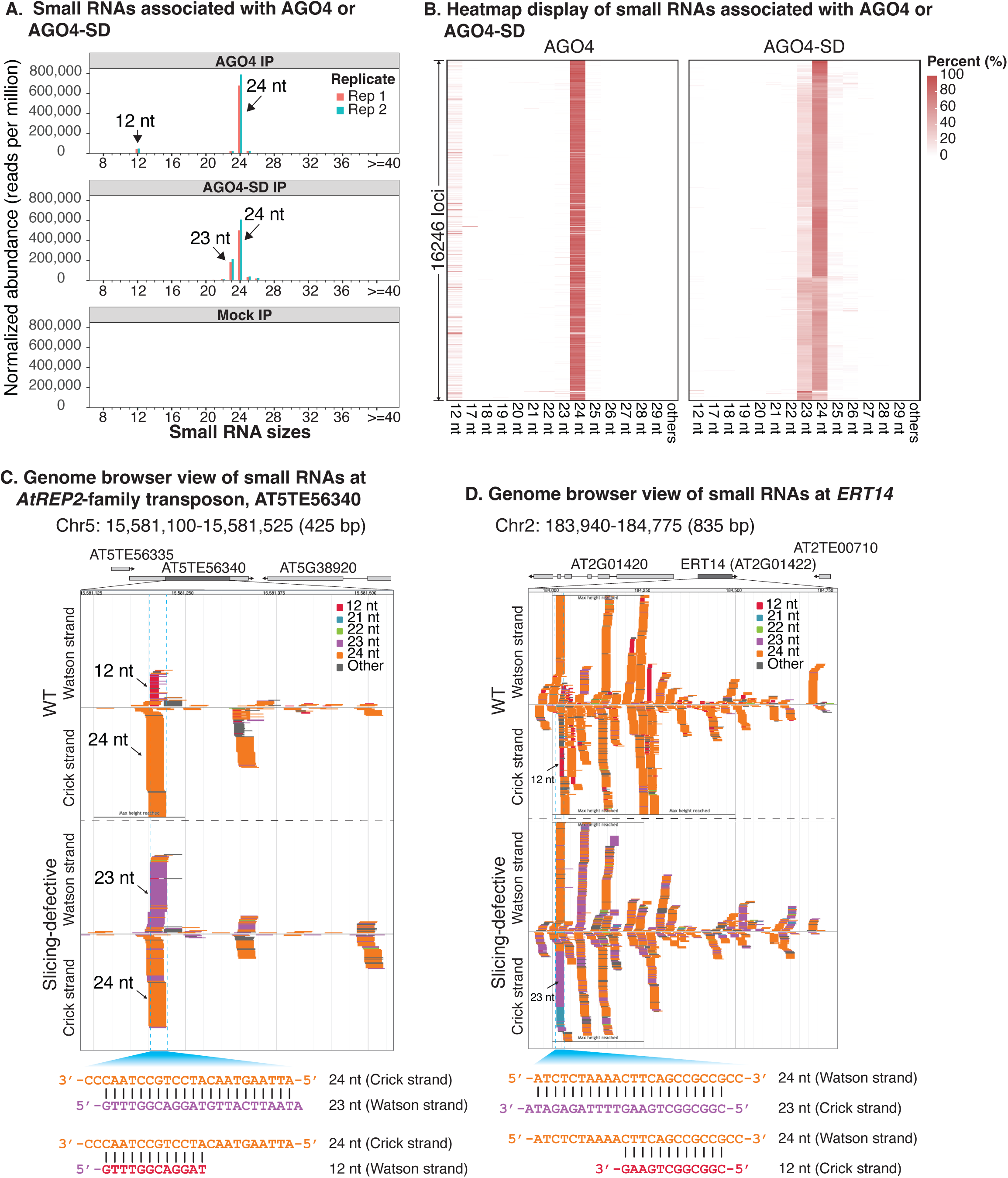
Distinct small RNA populations are associated with wild-type versus slicing-defective AGO4. **A**. Relative abundance of small RNAs associated with AGO4 or AGO4-SD. See also Supplementary Figures S1 and S2. **B**. Relative abundance of AGO4 or AGO4-SD associated RNAs of different size classes at 16,246 loci where 23 or 24 nt siRNAs are the dominant size class. **C and D**. Genome browser views of small RNAs copurifying with AGO4 or AGO4-SD. At5TE56340 (panel C) is an *AtREP2*-family transposon. ERT14 (panel D) is an RdDM locus characterized by (Blevins et al. 2014). RNAs are color-coded by size and displayed above or below the central horizontal axis according to strandedness. RNAs within the regions denoted by dashed lines were used for sequence alignments at the bottoms of panels C and D. See also Supplementary Figures S3 and S4.

Analysis of total small RNA abundance, without enrichment by AGO4 immunoprecipitation, revealed similar levels of 23 nt siRNAs in AGO4 and AGO4-SD plants (Figure S1C). Given that 23 nt RNAs appear to be efficiently sliced once 24/23 siRNA duplexes are loaded into AGO4 (see Figures. 1F and 2A), this finding, combined with prior RNA blot results showing that 23 and 24 nt siRNAs are equally sensitive to digestion by dsRNA-specific nuclease V1 and equally insensitive to ssRNA-specific RNase A (Blevins et al. 2015), suggests that most 23 nt siRNAs in cells exist within a pool of 24/23 duplexes that have not been loaded into AGO4.

Small RNAs that co-IP with both AGO4 and AGO4-SD have a strong signature for a 5’ terminal adenosine (A) among 24 nt siRNAs (Figure S2A). By contrast, the abundant 23 nt siRNAs that copurify with AGO4-SD have a strong signature for uridine (U), located one nucleotide upstream from the 3’ terminus (Figure S2B). This U occurs in the RDR2-transcribed RNA strand and is the complement of the 5’ terminal A of the Pol IV-transcribed 24 nt siRNA strand (Singh et al. 2019; Loffer et al. 2022). Its penultimate position is due to RDR2’s terminal transferase activity (Blevins et al. 2015) which adds an extra untemplated nucleotide onto the 3’ end of its transcripts, thus generating a 1 nt overhang relative to the Pol IV strand. Collectively, the 5’ A and penultimate 3’ U signatures are indicative of 24/23 siRNA duplexes that come from the end of the dsRNA precursor that includes the 5’ end of the Pol IV strand and the 3’ end of the RDR2 strand (Singh et al. 2019) (see Figure 1A).

RNAs of 22, 23, 25 or 26 nt, detected in trace amounts in association with wild-type AGO4, all have strong 5’ A signatures, just like 24 nt siRNAs (Figure S2A). Importantly, the 23 nt RNAs associated with wild-type AGO4 lack the penultimate 3’ U signal characteristic of unsliced 23 nt siRNAs associated with slicing-defective AGO4-SD (compare Figures S2A and S2B). Thus, the rare 23 nt siRNAs that associate with wild-type AGO4 are apparently 24 nt siRNAs that were truncated by 1 nt; they are not 23 nt passenger strands. Likewise, the 25 and 26 nt siRNAs found associated with both AGO4 and AGO4-SD appear to be 24 nt siRNAs elongated at their 3’ ends by untemplated uridylation (Figures S2B and S2C). Our detection of truncated and uridylated forms of 24 nt siRNAs that copurify with AGO4 suggests that siRNAs undergo a turnover process like miRNA turnover (Zhao et al. 2012).

### Sliced passenger strand fragments remain associated with AGO4

Approximately 16,200 loci give rise to the 24 and/or 23 nt siRNAs associated with AGO4 or AGO4-SD and account for >95% of the small RNAs detected by sequencing (Table S1). A heatmap display shows that 12 nt RNAs that co-IP with wild-type AGO4 (see Figure 2A, top panel), and 23 nt siRNAs that copurify with AGO4-SD, are broadly represented among these loci (Figure 2B).

Genome browser views of individual loci provided insight into the relationships among 12, 23 and 24 nt RNAs (Figures 2C and D). The 12 nt RNAs found associated with wild-type AGO4 come from the same strands as 23 nt RNAs that associate with slicing defective AGO4. In fact, manual alignment reveals that 12 and 23 nt RNAs share the same 5’ ends (see diagrams at the bottoms of Figures 2C and D). The 3’ termini of 12 nt RNAs occur at the positions where passenger strand slicing is predicted to occur, consistent with 12 nt RNAs being present in wild-type but not slicing defective AGO4 fractions. Collectively, these data suggest that 12 nt RNAs associated with AGO4 are the 5’ fragments of sliced passenger strand RNAs.

To extend our manual alignments of 23 and 24 nt RNAs at selected loci (e.g., those of Figures 2C and D) to all loci, and in an unbiased way, we conducted a computational prediction of strand pairing patterns among the 5,261,134 unique 24 or 23 nt RNA sequences associated with slicing-defective AGO4 (see Figure S3A and Supplemental Materials and Methods). The pairing configuration represented by the largest number of siRNA pairs has 23 nt siRNAs basepaired to 24 nt siRNAs such that the 5’ ends of the 24 nt strands are recessed by 1 nt relative to the 3’ ends of 23 nt strands (Figure S3B). No such arrangement was observed upon analysis of a computationally simulated library composed of small RNAs with the same abundance and sizes as the experimental dataset, and coming from same genomic loci, but initiating at random positions (Figures S3B and C).

24 nt RNAs outnumber 23 nt RNAs by approximately 3:1 in the pool of RNAs associated with slicing-defective AGO4 (Figure 2A), suggesting that 24/24 nt and 24/23 nt duplexes are loaded into AGO4 in similar abundance. Computational analysis of how 24 nt siRNAs can best pair with other, complementary 24 nt siRNAs among the pool of RNAs associated with AGO4-SD, predicts duplexes with 2 nt 3’ overhangs at each end as the most abundant duplex species (Figure S3C, left panel), which matches the known pairing of 24/24 duplexes generated by DCL3 dicing *in vitro* (Fukudome et al. 2021; Loffer et al. 2022). No such pattern was found for 24 nt RNAs associated with wild-type AGO4 (Figure S3C, right panel), consistent with the interpretation that slicing eliminates one strand of any 24/24 duplex that is loaded. It is noteworthy that regardless of whether a passenger strand is 23 or 24 nt in length, slicing will generate a 12 nt product due to cleavage opposite the guide strand’s 10^th^ nucleotide when measured from its 5’ end (see Figure S4).

### Passenger strand slicing can be recapitulated *in vitro* using recombinant AGO4

To enable biochemical assays of AGO4 slicing, we used baculovirus vectors to express N-terminal SUMO and FLAG-tagged wild-type or slicing-defective AGO4, with the latter having the catalytic triad mutated from DDH to AAA (abbreviated as AGO4-SD^AAA^)(Figure 3A), in insect cells. A 24 nt RNA was then incubated with the recombinant proteins to serve as a guide strand, followed by incubation with a complementary 23 nt siRNA, 5’ end-labeled with ^32^P (Figure 3B), yielding a duplex mimicking a DCL3-generated 24/23 duplex (Loffer et al. 2022)(see summary diagram in Figure S3B). In reactions involving wild-type AGO4, a labeled 12 nt RNA slicing product was generated (Figure 3B, lane 6), supporting the interpretation that 12 nt RNAs associated with wild-type AGO4 *in vivo* (Figure 2 and Figure S4) are slicing products. Recombinant AGO4-SD^AAA^ displayed no slicer activity, as expected (Figure 3B, lane 9).

**Figure 3.**
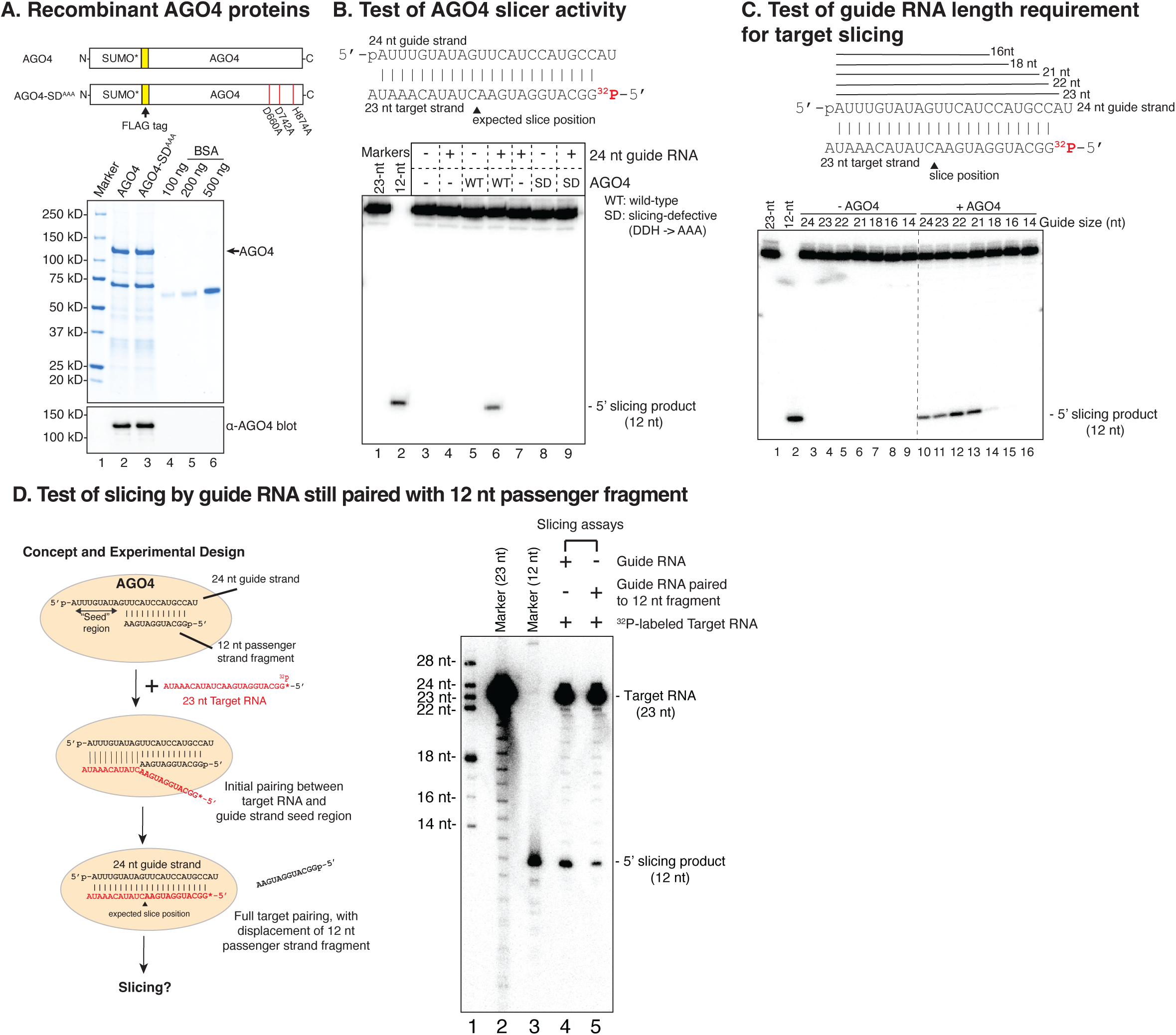
Guide strand-mediated slicing of 23 nt passenger strands can be recapitulated *in vitro* using recombinant AGO4. **A**. Diagrams of recombinant AGO4 and AGO4-SD^AAA^ in which aspartates D660 and D742 and histidine H874 were changed to alanines. A Coomassie-stained gel and anti-AGO4 immunoblot reveal similar expression levels for the two proteins. **B**. AGO4 has passenger strand slicer activity. The diagram shows the pairing of 24 nt and 23 nt strands of a DCL3-diced 24/23 duplex. The predicted cleavage site is denoted by a black triangle. In the experiment, AGO4 or AGO4-SD^AAA^ were first incubated with or without the 24 nt siRNA guide, then ^32^P end-labeled 23 nt siRNA was added. Slicing results in a 12 nt end-labeled fragment. RNAs were resolved by denaturing PAGE. End-labeled 23 nt and 12 nt size markers are present in lanes 1 and 2. Control reactions lacked added protein (lanes 3, 4 and 7) or guide RNA (lanes 3, 5 and 8). **C**. Guide RNAs of variable size program target RNA cleavage. Strands of 23, 22, 21, 18, 16 and 14 nt RNA that are 3’ truncations of the 24 nt guide strand shown are represented by solid lines. In the experiment, wild-type AGO4 was pre-incubated with guide strands of variable length then end-labeled 23 nt target RNA. End-labeled 23 nt and 12 nt size markers were run in lanes 1 and 2. AGO4 was omitted from reactions in lanes 3 to 9 and included in the reactions of lanes 10-16. See also Supplementary Figures S5 and S6. **D**. Test of slicing by a guide RNA paired with a 12 nt passenger strand fragment The cartoon at left illustrates the unpaired seed region of the guide RNA initiating pairing with a target RNA, leading to displacement of the 12 nt fragment and potential slicing of the target. In the autoradiogram at right, end-labeled size markers were resolved by denaturing PAGE in lanes 1 to 3. Recombinant AGO4 was loaded with the 24 nt single-stranded guide RNA (lane 4) or the guide RNA paired with the 12 nt passenger strand fragment (lane 5) and tested for the ability to slice a 5’ end-labeled 23 nt target RNA to produce a labeled 12 nt product.

We next tested the guide strand length requirements for AGO4 slicing. Interestingly, 21-24 nt guide RNAs, differing only at their 3’ ends, programmed equivalent levels of passenger strand slicing (Figure 3C, lanes 10 to 13). These results indicate that AGO4 does not anchor the guide RNA’s 3’ end at a fixed position within the PAZ domain, unlike human AGO2 and several other AGO proteins (Ma et al. 2004; Jinek and Doudna 2009; Frank et al. 2012). Examination of a predicted structure for AGO4, obtained using AlphaFold (Jumper et al. 2021), suggests that an extended b hairpin and disordered positively charged loop flanks the predicted 3’ end of the guide-strand RNA, in lieu of a nucleotide binding pocket, as in Human AGO2 and Arabidopsis AGO1 (Figure S5A). The sequences of the disordered loop are also conserved in the AGO4-related proteins AGO6 and AGO9 (Figure S5B; see also (Poulsen et al. 2013). We speculate that these extended motifs allow for siRNA 3’ ends of variable length.

A 12 nt passenger strand fragment basepaired to a 24 nt guide strand would leave the seed region of the guide strand (nucleotide positions 2-8, counting from the 5’ end) single-stranded and potentially still able to base pair with a target RNA (see the left panel in Figure 3D). To test this possibility, we annealed a 24 nt guide strand with an excess of a 12 nt RNA equivalent to a sliced passenger strand fragment, then incubated the resulting 24/12 nt duplex with recombinant AGO4. A 23 nt target RNA, 5’ end-labeled with ^32^P and complementary to the 24 nt guide RNA was then incubated with the complex. Slicing of the 23 nt RNA ensued, with AGO4 loaded with a 24/12 nt duplex displaying slicing activity comparable to AGO4 loaded only with a 24 nt guide RNA (Figure 3D, compare lanes 4 and 5).

### Guide strand end-modifications and terminal nucleotide requirements

Results of previous studies suggested that Pol IV-transcribed strands of siRNA precursor dsRNAs have 5’ monophosphates (Blevins et al. 2015; Li et al. 2015; Zhai et al. 2015) whereas RDR2-transcribed strands have 5’ triphosphates (Singh et al. 2019) (see Figure 4A). These end groups persist in diced siRNAs (Singh et al. 2019). To test whether 5’ end modifications affect guide strand loading or activity, we incubated FLAG-tagged recombinant AGO4 with 24 nt RNAs having either 5’ hydroxyl, 5’ monophosphate or 5’ triphosphate groups. Following affinity capture of AGO4 using anti-FLAG Dynabeads, associated RNAs were subjected to RNA blot analysis. The results show that guide RNAs can be loaded into AGO4 regardless of their 5’ end modifications, but the strongest signals were observed for RNAs with a 5’ monophosphate group (Figure 4B, lane 6 compared to lanes 7 and 8). Guide RNAs with 5’ monophosphate groups also programmed 5 to 8-fold higher levels of target slicing than guide RNAs with 5’ hydroxyl or triphosphate groups (Figure 4C, lane 6 compared to lanes 7 and 8). The observed 5’ monophosphate preference suggests that 24 nt siRNAs derived from the 5’ ends of Pol IV strands may be preferentially loaded into AGO4 and may preferentially program slicing compared to siRNAs derived from the 5’ triphosphorylated RDR2 strand. MID domain amino acids known to be involved in 5’ phosphate interactions in Arabidopsis AGO1 are highly conserved, in sequence and spatial position, in AGO4 (Figures S6A and B) and all Arabidopsis AGO proteins (Figure S6B).

**Figure 4.**
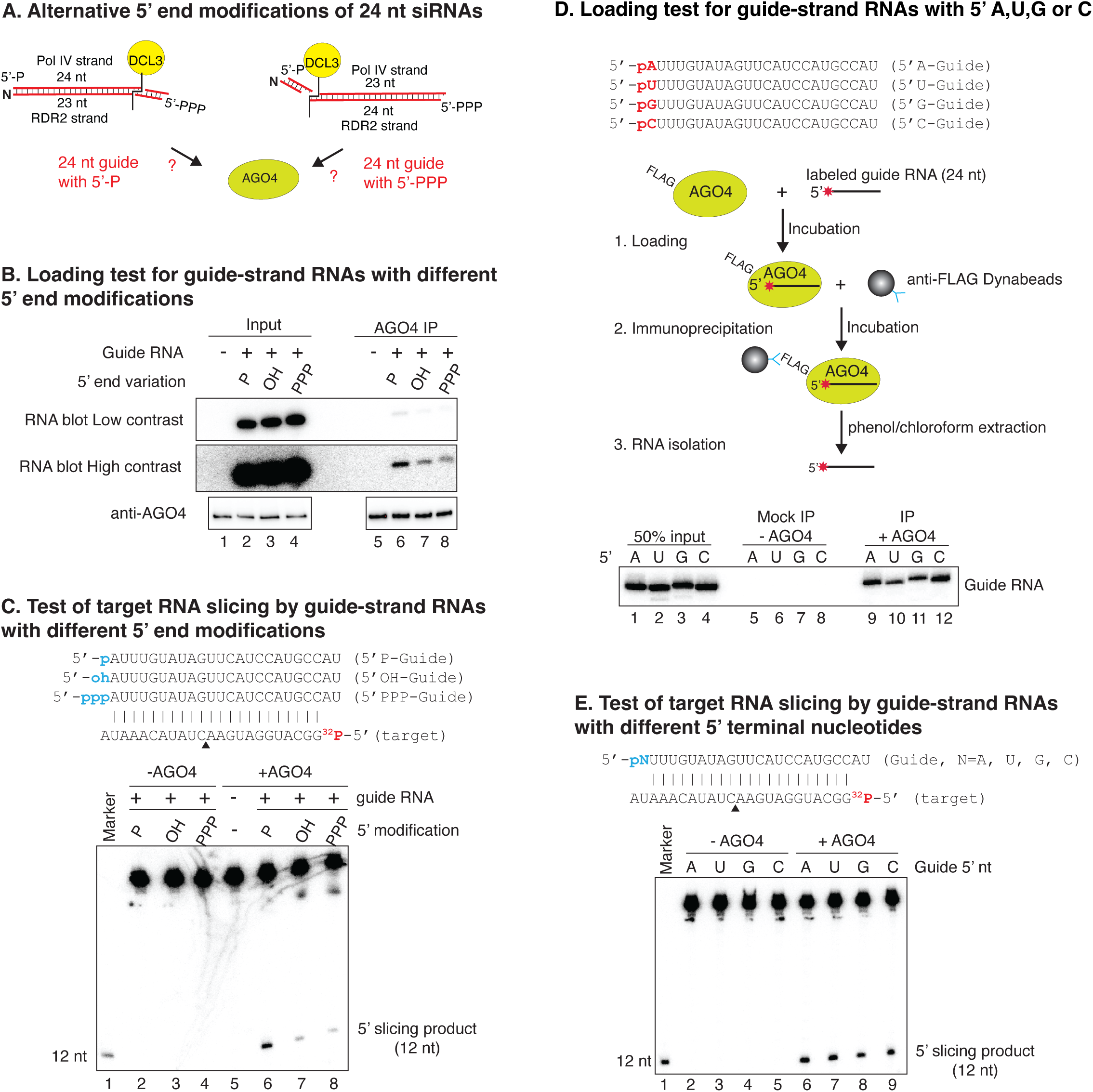
Tests of guide strand 5’ chemical groups and nucleotides in AGO4 loading and slicing. **A**. Cartoon illustrating alternative features of precursors and diced siRNA duplexes. **B**. Test of AGO4 loading for 24 nt guide RNAs with 5’ monophosphate (P), hydroxyl (OH), or triphosphate (PPP) groups. Input samples are shown in lanes 1-4. RNAs that co-IP with AGO4 are in lanes 5-8. RNAs were subjected to denaturing PAGE, transferred to membranes, and detected by RNA blot hybridization. Controls in lanes 1 and 5 lacked RNA. **C**. Target RNA slicing programmed by guide siRNAs with different 5’ end modifications. 24 nt guide RNAs with different 5’ groups, paired with a 5’ end-labeled 23 nt target RNA, are shown at the top. The predicted slice site is denoted by a black triangle. The alternative guide RNAs were incubated with or without AGO4 followed by incubation with ^32^P-labeled 23 nt target RNA. RNAs were then subjected to denaturing PAGE. An end-labeled 12 nt size marker was run in lane 1. Lane 5 is a control in which AGO4 was included but guide RNA was omitted. **D.** Test for preferential loading of guide RNAs with different 5’ nucleotides. 24 nt RNAs with each of the four possible nucleotides at their 5’ ends were ^32^P-end-labeled and incubated with FLAG-tagged recombinant AGO4. RNAs associated with AGO4 captured on anti-FLAG Dynabeads were resolved by denaturing PAGE (lanes 9-12). In mock IP control reactions (lanes 5-8), AGO4 was omitted. Lanes 1 to 4 show input RNA. See also Supplementary Figure S7. **E**. Test of target RNA slicing programmed by guide RNAs with different 5’ nucleotides. The different guide RNAs were incubated with (lanes 6 to 9) or without AGO4 (lanes 2 to 5) then incubated with end-labeled 23 nt target RNA. RNAs were then purified, subjected to denaturing PAGE and autoradiography.

Approximately 80% of the 24 nt siRNAs associated with AGO4 *in vivo* begin with adenosine, suggesting that AGO4 may selectively bind 24 nt siRNAs with a 5’ A. As a test of this hypothesis, we synthesized 24 nt RNAs that differ only by having 5’ A, U, C or G. We then end-labeled the RNAs with ^32^P, incubated them with FLAG-tagged recombinant AGO4, affinity captured the resulting AGO4-RNA complexes and subjected the RNAs to denaturing PAGE and autoradiography. This experiment revealed that 5’ nucleotide identity has little effect on AGO4 binding (Figure 4D, lanes 9 to 12). We also performed a competition assay in which AGO4’s binding of a ^32^P-labeled 24 nt RNA with a 5’ A was challenged by inclusion of increasing amounts of phosphorylated but unlabeled 24 nt siRNAs that have A, U, G, or C nucleotides at their 5’ ends. All four competitor RNAs competed effectively against the labelled RNA (Figure S7). Likewise, guide RNAs with 5’ A, U, C or G nucleotides programmed similar levels of target RNA slicing (Figure 4E). Collectively, the data of Figures 4 and S7 indicate that 5’ terminal nucleotide identity has little effect on AGO4 guide strand loading or enzymatic activity.

### AGO4 retains sliced target RNA fragments *in vitro* and *in vivo*

Our detection of 12 nt passenger strand fragments in association with AGO4 led us to test the extent to which sliced RNA fragments are released or retained by AGO4 (Figure 5). For these experiments, FLAG-tagged AGO4 expressed in transgenic plants was immobilized on anti-FLAG Dynabeads and loaded with a 24 nt RNA guide strand. A complimentary 5’ end-labelled target RNA was then added. After incubation, reactions were separated into supernatant and bead-associated fractions, with the bead fractions subjected to multiple rounds of washing. RNAs of both fractions were then purified, resolved by denaturing PAGE, and visualized by autoradiography (Figure 5A). Using a 23 nt target RNA, mimicking a passenger strand, an end-labeled slicing product of 12 nt was detected primarily in the washed bead fraction, not the supernatant (Figure 5B, compare lanes 4 and 5). Changing guide strand nucleotides 9-11 to make them non-complementary to the corresponding 23 nt target RNA nucleotides impaired slicing, demonstrating the importance of basepairing near the cut site (Figure 5B, compare lanes 6-7 to lanes 4-5). Next, we tested AGO4’s ability to slice a 51 nt target RNA end-labeled with ^32^P (Figure 5C), yielding a labeled 29 nt fragment that remained associated with bead-immobilized AGO4 (Figure 5C, lane 5 compared to lane 4).

**Figure 5.**
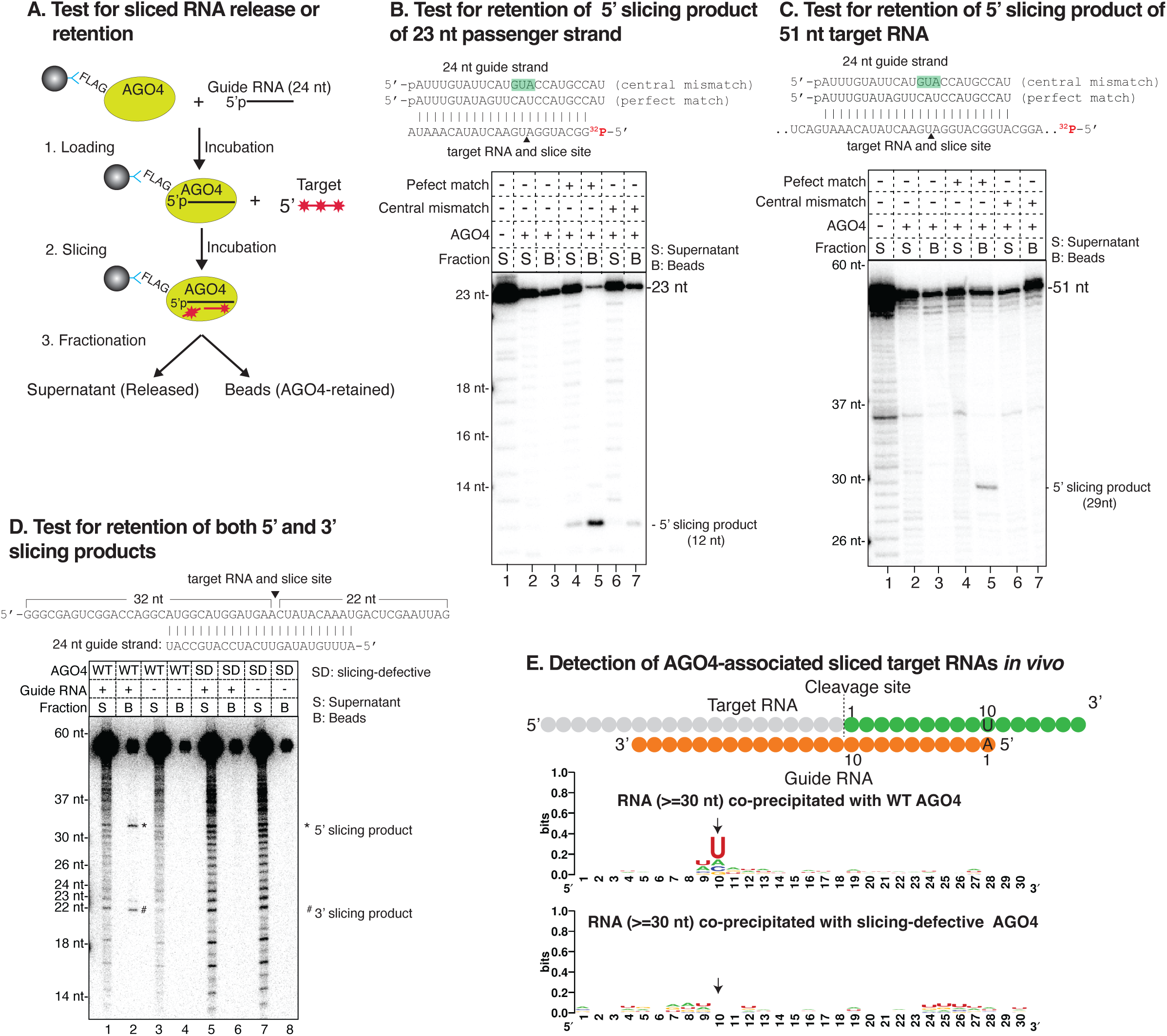
AGO4 retains sliced target RNAs *in vitro* and *in vivo*. **A**. Cartoon of the strategy for testing AGO4 release or retention of sliced target RNAs. **B**. Test for retention of 23 nt target strand 5’ cleavage products. At the top are sequences of a 24 nt guide RNA with 100% complementarity to the ^32^P-labeled target RNAs and a guide RNA with three nucleotides of target non-complementarity (highlighted in green). In the experiment, FLAG-AGO4 immobilized on anti-FLAG Dynabeads was incubated with buffer only (lanes 2 and 3), with the 24 nt guide RNA having perfect complementarity to the target RNA (lanes 4 and 5), or with 24 nt guide RNAs with mismatches to the target RNA (lanes 6 and 7). RNAs in the supernatant (S) and washed bead (B) fractions were purified, subjected to denaturing PAGE and autoradiography. Lane 1 is an anti-FLAG IP control using non-transgenic plants. **C**. Test for retention of 5’ cleavage products of a 51 nt RNA target. The experiment was conducted as for panel B. **D**. Both 5’ and 3’ cleavage products of long RNAs are retained by AGO4 *in vitro*. The experiment was conducted as for panel B except that a body-labeled 54 nt target RNA was used. The expected sizes of its slicing products are 22 and 32 nt. **E**. Sliced products of RdDM target locus RNAs are retained by AGO4 *in vivo*. The cartoon shows how 24 nt guide RNAs of variable sequence (orange) would pair with complementary target RNAs. 5’ and 3’ cleavage products are color-coded grey and green respectively. At position 10 of the 3’ slicing products (green fragments), a strong uridine (U) signature is expected due to complementarity to the adenosine (A) at the guide strand 5’ end. The sequence logos below show RNA-seq analyses for RNAs longer than 30 nt that co-IPed with wild-type AGO4 or AGO4-SD. The arrow denotes the position of the expected U signature.

Using 5’ end-labeled target RNAs only allows 5’ slicing products to be detected. This led us to test a body-labeled 54 nt target RNA generated via incorporation of an alpha-^32^P-labeled nucleotide during transcription by T7 RNA polymerase *in vitro*. Labeled fragments of 32 nt and 22 nt were generated upon AGO4 slicing, corresponding to both the 5’ and 3’ cleavage products (Figure 5D, lane 2), and both were enriched in the AGO4-bead fraction. This experiment indicates that both 5’ and 3’ slicing products remain associated with AGO4 after target RNA cleavage. Control experiments confirmed that slicing products are not generated in the absence of a guide RNA (Figure 5D, lane 4) or by slicing-defective AGO4 (Figure 5D, lanes 6 and 8).

Our finding that AGO4 retains its slicing products *in vitro* prompted us to ask whether scaffold RNAs at RdDM loci, primarily synthesized by RNA polymerase V (Wierzbicki et al. 2009), are also sliced and retained *in vivo*. A prediction is that sliced scaffold RNAs would include RNAs with a strong signature for a uridine (U) at position 10. This is because AGO4-loaded 24 nt guide RNAs have an ∼80% bias for a 5’ terminal adenosine (A) and because slicing occurs opposite the 10^th^ position measured from this 5’ A. Thus, the expected U at position 10 in of a sliced target RNA would be the complement of the guide strand’s 5’ terminal A (Figure 5E, see the model in the top panel). With this prediction in mind, we analyzed the sequences of all RNAs that co-immunoprecipitated with wild-type AGO4 *in vivo*, examining RNAs of 30 nt or longer. A strong signature for U is indeed present at position 10 of these AGO-associated RNAs (Figure 5E). No +10 U signature was observed for RNAs associated with slicing-defective AGO4-SD (Figure 5E).

### AGO4 slicer activity elevates DNA methylation levels at RdDM loci, genome-wide

We tested the importance of AGO4’s slicing ability for DNA methylation at RdDM loci using whole-genome bisulfite sequencing. In *ago4-4* null mutant plants, compared to wild-type plants, cytosine methylation at CHH and CHG motifs is substantially reduced at 6,718 genomic loci whereas CG methylation is mostly unaffected (Figure 6A, compare left three panels). We define these 6,718 loci as AGO4-dependent differentially methylated regions (agoDMRs; see Supplemental Table 2). Rescue of the *ago4-4* mutant by the wild-type AGO4 transgene restored CHG and CHH methylation to nearly wild-type levels. In contrast, expression of AGO4-SD resulted in CHG and CHH methylation levels intermediate between those of the mutant and wild-type, and this was true for the majority of agoDMR loci (see heatmap panel of Figure 6A). AGO4-dependent CHG and CHH methylation is particularly apparent at short TEs (1-2 kb) and the edges of long TEs (> 4kb) (Figure S8), consistent with RdDM mainly targeting these regions (Zemach et al. 2013).

**Figure 6.**
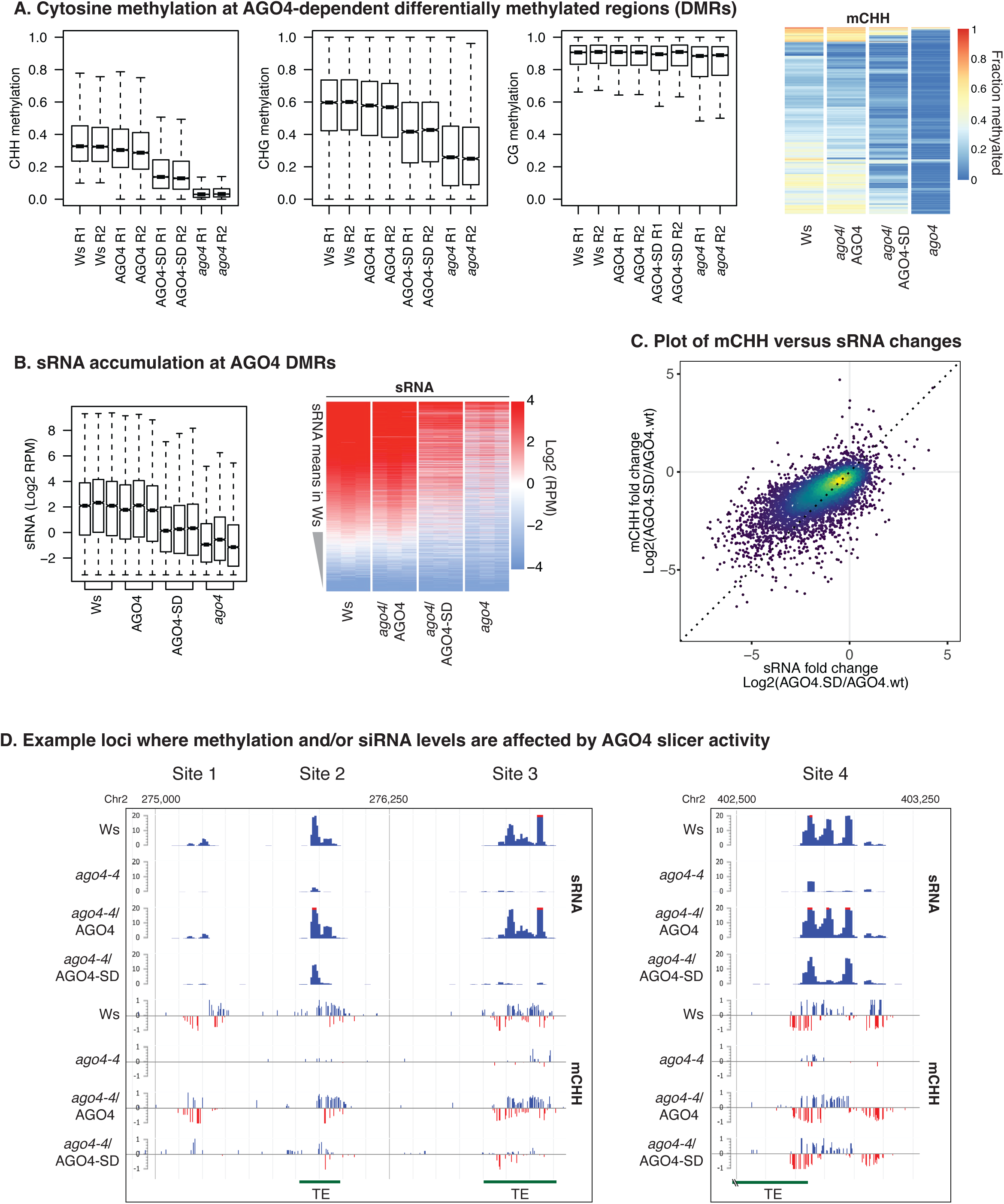
AGO4 slicer activity enhances non-CG methylation at virtually all RdDM loci. **A**. Cytosine methylation at AGO4-dependent differentially methylated regions (DMRs). **B**. Relative small RNA abundance at AGO4 DMRs displayed as boxplots and as a heatmap. siRNA accumulation was normalized to the total mappable read count of the corresponding small RNA library (RPM: read per million). **C**. Correlation between CHH methylation and siRNA levels at AGO4 DMRs. Points overlapping at higher density are shown in warmer colors. **D**. Representative genome browser views of loci at which *de novo* DNA methylation and/or small RNA accumulation is affected by AGO4 slicer activity. See also Supplementary Figure S8.

**Figure 7.**
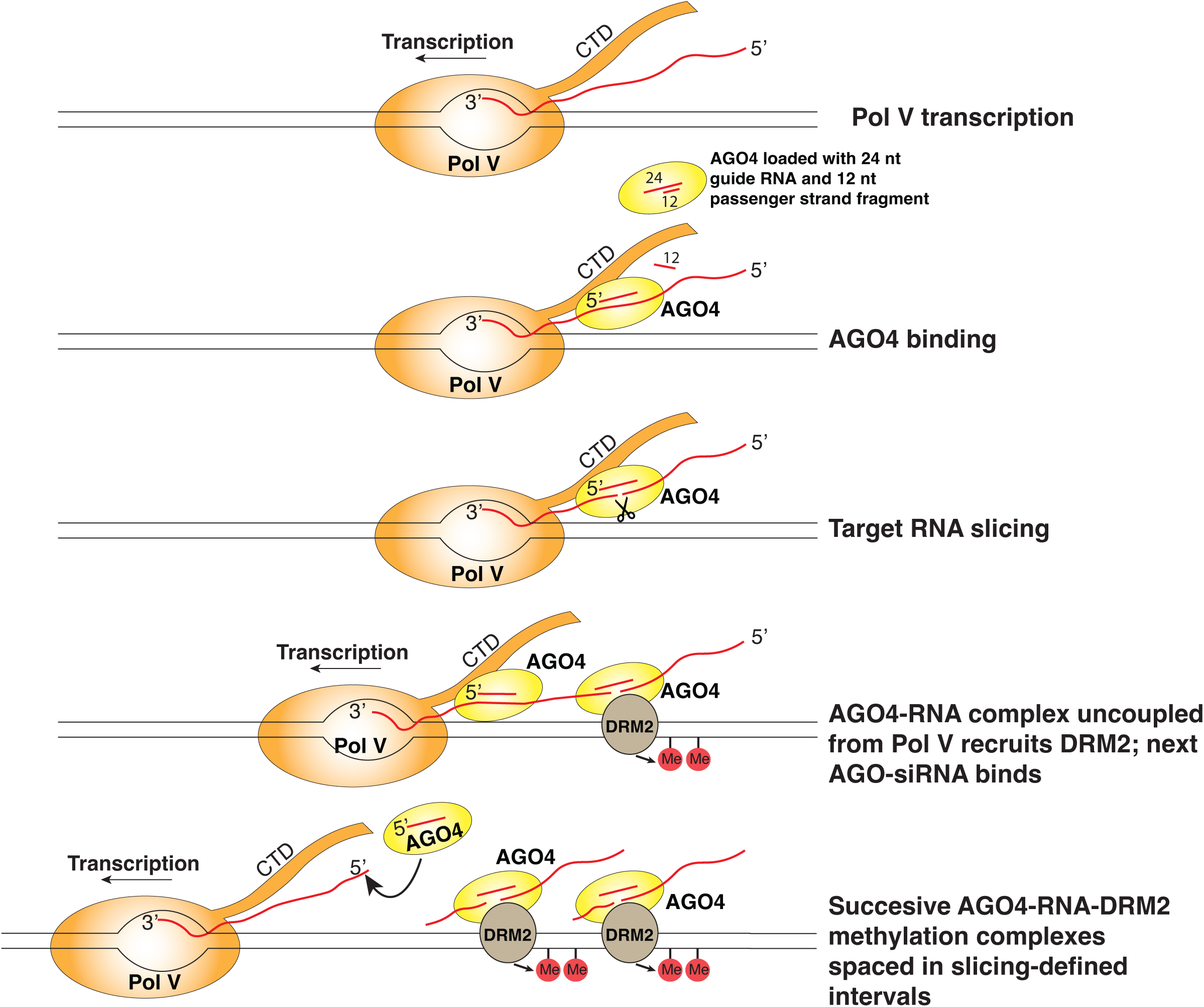
A model for discontinuous RNA-directed DNA methylation in intervals reflecting AGO4 slicing.

Analyses of siRNA levels by RNA-seq revealed that in the *ago4-4* mutant, siRNA levels at agoDMR loci decrease, paralleling the decreases in CHG and CHH methylation levels. Like cytosine methylation, siRNA levels are nearly fully restored by wild-type AGO4 but are only partially restored byAGO4-SD (Figure 6B). The positive correlation between CHH methylation levels and small RNA levels is illustrated in Figure 6C.

The data of Figures 6A, B, and S8 suggest that AGO4 slicing has primarily quantitative, rather than qualitative, effects on siRNA and CHH methylation levels at agoDMR loci. However, within individual loci, differences in the spatial distribution of siRNAs and CHH methylation are often apparent, as shown in genome browser views for four loci (Figure 6D). At sites 1 and 3, CHH methylation levels are severely impacted in the *ago4-4* mutant and are restored to wild-type levels by the wild-type AGO4 transgene but are only weakly rescued by the AGO4-SD transgene, which also fails to restore siRNA levels. In comparison, sites 2 and 4 have two or three spatially separated peaks of siRNAs that are greatly diminished in the *ago4-4* mutant and are restored by wild-type AGO4. At each of these loci, the AGO4-SD protein fails to restore one of the siRNA peaks, which correlates with reduced methylation within that interval (most apparent for site 2). Collectively, these results indicate that quantitative differences in slicing-dependent methylation and siRNA levels at RdDM loci can frequently be attributed to differences within sub-intervals of affected loci.

## Discussion

### The function of 23 nt siRNAs revealed

DCL3-generated 23 nt RNAs derive from the 3’ ends of Pol IV or RDR2 strands of siRNA precursors and are basepaired to 24 nt siRNAs derived from precursor 5’ ends (Singh et al. 2019; Loffer et al. 2022), but whether 23 nt RNAs play a role in RdDM has long been a question. Our results provide an answer, showing that 23 nt RNAs function as passenger strands for 24 nt siRNA guide strands. The 24 nt guide strands then program AGO4 to slice the 23 nt passenger strands with nearly 100% efficiency. What few 23 nt RNAs remain associated with AGO4 lack the 3’ penultimate U signature of passenger strands; instead, they have sequences indicative of truncated 24 nt strands (see Figure S2). Collectively, our results indicate that 23 nt siRNAs do function in RdDM, but only indirectly - by specifying that their 24 nt partners are loaded into AGO4 to carry out subsequent target locus interactions.

### Old and new hypotheses for AGO4 guide strand selection

How is the 24 nt strand of a 24/23 duplex selected as the guide strand? A longstanding hypothesis has been that AGO4 has a structural requirement for 24 nt RNAs, such that 23 nt RNAs cannot be stably bound. However, our biochemical results show that AGO4 can be loaded with 21, 22, 23 or 24 nt RNAs, all of which guide target RNA slicing with similar efficiency (see Figure 3). Intriguingly, low-abundance Pol IV-dependent 21-22 nt siRNAs are generated at loci where 24 nt siRNAs accumulate and can participate in RdDM (Panda et al. 2020). Our biochemical results support the hypothesis that these 21-22 nt siRNAs could potentially be loaded into AGO4, as is the case for AGO6, which incorporates 21-22 nt TE or RNAi-derived siRNAs and guides DNA methylation in the non-canonical RdDM pathway (McCue et al. 2015).

It has also been thought that AGO4 may actively select guide strands with a 5’ terminal adenosine, because ∼80% of 24 nt siRNAs associated with AGO4 have 5’ adenosines (Mi et al., 2008). Precedence for this idea includes structural studies of human AGO2, whose nucleotide specificity loop in the MID domain favors binding of a 5’ U or A (Frank et al. 2010; Frank et al. 2012) and studies of *Arabidopsis* AGO1, whose similar specificity loop favors binding of miRNAs with a 5’ U (Mi et al. 2008; Frank et al. 2012). However, our results show that 24 nt siRNAs with 5’ A, U, C or G can associate with AGO4 and mediate target RNA cleavage with similar efficiencies (see Figures 5D and E). Taken together, our tests of guide strand length and terminal nucleotide identity lead us to conclude that AGO4 binds the siRNAs available to it, as dictated by the activities of the enzymes that generate the siRNAs, namely Pol IV, RDR2 and DCL3 (Singh et al. 2019). Pol IV most frequently initiates transcription with an A or G; likewise, RDR2 has a (weaker) tendency to initiate with A or U (Singh et al. 2019). Thus, enrichment for 5’ adenosines is a feature of siRNA precursors. DCL3 then displays a preference for binding dsRNA ends that include a 5’ A, and discriminates against 5’ G, and then measures and cuts 24 nt downstream, resulting in biased production of 24 nt siRNAs with a 5’ A (Loffer et al. 2022).

However, the question of why 24 nt, and not 23 nt RNAs of 24/23 duplexes are selected as guide strands remains. The asymmetry inherent to 24/23 duplexes is likely important, with 3’ ends of 23 nt strands overhanging by 1 nt and 3’ ends of the 24 nt strands overhanging by 2 nt. We speculate that this asymmetry is recognized by an as-yet unidentified AGO4 loading activity that binds and orients the duplex such that the 24 nt strand is stably incorporated into AGO4. In the case of 24/24 duplexes that are also generated by DCL3, and composed of siRNAs from the 3’ ends of Pol IV strands and the 5’ ends of RDR2 strands (Loffer et al. 2022), 3’ overhangs of 2 nt are present at each end, such that these duplexes are symmetrical. If a 3’ overhang of 2 nt is what identifies the strand to be loaded, the 24 nt strand of a 24/23 duplex would be chosen, and either strand of a 24/24 duplex could be chosen.

### Retention of sliced RNAs suggests a model for how slicing may enhance RdDM

Our finding that 12 nt 5’ fragments of sliced 23 nt passenger strands remained associated with AGO4, both *in vivo* and *in vitro* (see Figures 2 and 5B) was unexpected. Presumably the 12 nt fragments are basepaired to the 3’ half of 24 nt guide strands, leaving the seed region of the guide strands single-stranded and available to initiate pairing with complementary target RNAs. Our test of this hypothesis showed that a 24 nt guide strand -12 nt passenger strand fragment duplex can, indeed program target RNA slicing (see Figure 3D). We speculate that the 12 nt fragment is displaced upon target RNA binding given that slicing requires perfect guide strand-target RNA basepairing surrounding the slice site (see Figure 5, panels B and C) and the 12 nt RNA overlaps this region.

An unanswered question is whether the 12 nt passenger strand slicing product is important for some aspect of AGO4 function. One possibility is that it helps protect the 3’ end of the paired 24 nt guide RNA and may be accommodated by the AGO4 PAZ domain’s predicted extended hairpin and disordered loop, which differs from the end-tethering pockets of other AGOs (see Figure S5). Another possibility is that a 12 nt passenger strand fragment basepaired to a guide strand serves as a mark of a new and unused AGO4-siRNA complex that has not yet seen action by engaging a complementary target RNA. Such a mark could potentially play a role in subcellular trafficking or docking of AGO4-siRNA complexes with Pol V or other components of the RdDM machinery.

A pioneering study indicated that AGO4 slicing competence affected methylation and siRNA abundance at some, but not all, of a limited set of Arabidopsis RdDM loci known at the time (Qi et al., 2006). Revisiting the question using bisulfite sequencing and small RNA deep sequencing to achieve a whole genome analysis shows that AGO4 slicer activity is needed to achieve maximal DNA methylation and siRNA levels at nearly all RdDM loci, not just a specific subset of loci (Figures 6A, B, and S8). But what does AGO4 slice? Early studies showed that AGO4 can be crosslinked to Pol V transcripts and that Pol V transcript levels detected by RT-PCR are higher in *ago4* mutants (Wierzbicki et al. 2009). Thus, Pol V transcripts have long been assumed to be likely targets of AGO4 slicing. Moreover, RNA sequencing of nascent Pol V transcripts generated under nuclear run-on conditions revealed a strong +10 U signature that, together with genetic evidence, suggested co-transcriptional slicing by one or more members of the AGO4 clade (Liu et al., 2018). Our study adds to these previous results by showing that AGO4 can account for the +10 U signature and that AGO4 surprisingly remains associated with the sliced products of nascent Pol V transcripts.

Why should AGO4 retain its slicing products? Importantly, Pol V transcripts are thought to be longer than 100 nt (Bohmdorfer et al. 2016) and they overlap genomic positions at which 24 nt siRNAs are generated in swarms (Wierzbicki et al. 2012), beginning and ending at variable positions throughout the loci (see Figures 2C and 2D). Thus, multiple 24 siRNAs can potentially basepair with any given Pol V transcript. There is also compelling evidence that protein-protein interactions occur between the C-terminal domain of the Pol V largest subunit and AGO4 (El-Shami et al. 2007), likely in a complex also involving AGO4 interactions with SPT5L (Lahmy et al. 2016; Liu et al. 2018). If AGO4-CTD interactions and siRNA-Pol V transcript interactions occur simultaneously, and co-transcriptionally, can these interactions be maintained as Pol V transcription elongation proceeds and the distance (measured in RNA length) between the polymerase and an AGO4-siRNA complexes progressively increases? Based on these considerations, we propose a model, shown in Figure 7, in which protein-protein interactions between the Pol V CTD and AGO4 initially occur coincident with siRNA-Pol V transcript basepairing. AGO4 slicing of the Pol V transcript ensues, allowing the 5’ portion of the nascent transcript to be uncoupled from the still-elongating polymerase, and with the AGO4-siRNA complex remaining bound to the sliced and released RNA. This process might then be repeated by the binding, slicing and release of successive AGO4-siRNA complexes as Pol V transcription proceeds. A further speculation is that the sliced RNA fragments retained by AGO4 may facilitate direct tethering at the corresponding DNA loci, perhaps contributing to evidence for R-loops at RdDM loci (Xu et al. 2017). Mass spectrometry evidence indicates that AGO4 and the *de novo* DNA methyltransferase, DRM2 can associate (Zhong et al. 2014). Thus, individual tethered AGO4-RNA complexes may be sufficient for DRM2 recruitment and methylation of corresponding DNA intervals, with methylation able to proceed at a pace independent of the rate of Pol V transcriptional elongation thanks to AGO4 slicing. Our finding that cytosine methylation and associated siRNA production can differ within RdDM locus sub-intervals that can also differ in their dependence on AGO4 slicing ability (see Figure 6) is compatible with the hypothesis that multiple AGO4-DRM complexes may act in parallel, in intervals at least partially defined by AGO4 slicing. Testing the model of Figure 7 will be a challenge for future studies.

## Materials and Methods

### AGO4 protein expression and purification

AGO4 proteins used in this study were expressed in *A. thaliana* or insect cells and purified as detailed in Supplemental Materials and Methods. Purified AGO4 proteins were resolved in 4-20% Mini-PROTEAN stain-free gels (Bio-Rad) and visualized by Coomassie Blue staining or immunoblotting with HRP-conjugated anti-FLAG M2 antibodies (Sigma) or anti-AGO4 polyclonal antibodies (Pikaard lab antibody stock #: 18187).

### Analysis of AGO4-bound RNAs

RNAs copurifying with wild-type AGO4 or slicing-defective AGO4-SD from *A. thaliana* were either 5’ end-labeled by [γ-^32^P] ATP and resolved in polyacrylamide gels under denaturing condition or subjected to high-throughput RNA sequencing. Detailed methods for RNA preparation can be found in Supplemental Materials and Methods.

### RNA oligonucleotide preparation

RNA oligos (Supplemental Table S3) were synthesized by Integrated DNA Technologies (IDT), except for the triphosphorylated oligo used in Figures. 4B and C, which was synthesized by BioSynthesis, Inc. RNA oligos were gel-purified (Nilsen 2013)). Purified RNA oligos were either phosphorylated with unlabeled ATP or 5’ end-labeled using [γ-^32^P] ATP (Perkin Elmer) and T4-Polynucleotide kinase (New England Biolabs), followed by purification using Performa spin columns (EdgeBio). ^32^P-labeled RNA oligos were gel-purified again to remove degraded RNAs.

### *In vitro* guide RNA loading and target RNA slicing assay

Immunoprecipitated AGO4 from transgenic *A. thaliana* or partially purified AGO4 from insect cells (detailed in Supplemental Materials and Methods) were incubated with 40 nM of unlabeled phosphorylated guide RNA in 20 µL of reaction buffer (100 mM Potassium acetate, 25 mM HEPES-KOH, pH 7.4, 5 mM Magnesium acetate, 10% glycerol, 0.01% IGEPAL, 0.5 mM DTT and 1 mM PMSF) for 1 hour at room temperature to allow guide RNA incorporation into AGO4. 1 µL ^32^P-phosphorylated target RNA (∼1000 cpm/µL) and 1 µg yeast tRNA (Thermo Fisher Scientific) was added to each reaction. After 1 hour incubation at room temperature, reactions were quenched by adding 200 µL of stop solution (300 mM NaOAc and 7M urea). RNAs were extracted by 200 µL of phenol/chloroform/isoamyl alcohol, ethanol precipitated, resuspended in 1X formamide loading buffer (40% deionized formamide, 0.5 mg/mL xylene cyanol, 0.5 mg/ml bromophenol blue, and 5 mM EDTA, pH 8.0), resolved in 15% polyacrylamide gels containing 7M urea, and visualized by phosphorimaging.

### Small RNA sequencing and data analyses

RNAs co-immunoprecipitating with AGO4 in inflorescence tissues were purified and used to generate small RNA sequencing libraries using Illumina’s TruSeq Small RNA Library Prep Kit. The cDNA libraries were single-end sequenced using an Illumina NextSeq500 platform. The 3’ adapter sequences were removed using Cutadapt v1.18 (Martin 2011) with the following options: *-a TGGAATTC --discard-untrimmed -e 0 -m 8 -O 8*. Trimmed reads were aligned to the *A. thaliana* TAIR10 genome using ShortStack v3.8 (Johnson et al. 2016) allowing no more than 2 mismatches. The alignment files were loaded in JBrowse configured with ‘Small RNA plugin’ (Hofmeister and Schmitz 2018) with modifications to allow customized small RNA alignment visualization. Sequence logos were prepared using WebLogo 3.6 (Crooks et al. 2004). Heatmaps for siRNA loci where 24 nt siRNAs are the predominant class were clustered by complete linkage with Euclidean distances calculated. Detailed methods for in silico small RNA duplex reconstitution can be found in Supplemental Materials and Methods.

### Whole-genome bisulfite sequencing and methylome data analysis

Genomic DNA was extracted from inflorescence tissues using a Nucleon Phytopure DNA Extraction Kit (Cytiva). Whole-genome bisulfite sequencing libraries for two biological replicates of each genotype were prepared using Perkin Elmer’s Nextflex Bisulfite-Seq Kit with approximately 500 ng of fragmented genomic DNA. Unmethylated cytosines in the adapter-ligated DNA were chemically converted by EZ DNA Methylation-Gold Kit (Zymo Research Corp.). The CT converted products were purified and subjected to 12 cycles of PCR amplification before single-end sequencing using an Illumina NextSeq500 platform.

To analyze the whole-genome bisulfite sequencing data, 3’ adapter sequences were first removed by Cutadapt v1.18 (Martin, 2011) with the following options: *-aAGATCGGAAGAGCACACGTCTGAACTCCAGTCAC -m 15 -q 20*. Trimmed reads were aligned to the *Arabidopsis* TAIR10 genome using bsmap2 (Xi and Li 2009) with the following options: *-w 100 -n 1 -p 3 -v 0.08*. The *Arabidopsis* genome sequence was binned into 100 nt windows with CG, CHG, and CHH (where H represents any nucleotide except G) methylation rate calculated for each window. To avoid regions with low sequencing coverage, only bins with at least 4 cytosines that were each covered by no less than 4 reads were included in the analysis. Bins that had at least 10% less CHH methylation in two independent replicates of *ago4-4* compared to two independent replicates of wild-type plants were defined as differentially methylated bins. Differentially methylated bins within 200 bp from each other were then merged into differentially methylated regions (DMRs).

## Data availability

AGO4-immunoprecipitated RNA sequencing and whole genome bisulfite sequencing data sets are deposited at GEO with accession number GSE201235. In-house scripts for these analyses are available at: https://github.com/wangfeng3392/small_RNA.

## Additional materials and methods

Additional materials and methods can be found in Supplemental Materials and Methods.

## Acknowledgements

We thank Michael Axtell for sharing *ago4-4*, *ago4-4/FLAG-AGO4* and *ago4-4/FLAG-AGO4-SD* seeds, Tony Yu and Zheng Tian for help collecting AGO4 transgenic plant tissues, and Ramya Enganti for repeating some *in vitro* slicing assays. We thank the Center for Genomics and Bioinformatics at Indiana University Bloomington (IUB) for library preparation and high-throughput sequencing, the Drosophila Genome Resource Center at IUB for insect cell culture facilities, and the IUB National Center for Genome Analysis Support (NCGAS) and University Information Technology Services for providing supercomputing resources. This research was supported by NIH grant GM077590 and funds to CSP as an Investigator of the Howard Hughes Medical Institute. JS was supported, in part, by a Carlos O. Miller graduate fellowship at IUB. HYH was supported by a Ruth L. Kirschstein National Research Service Award (NRSA).

## Author contributions

FW, HYH, and CSP designed the experiments. FW performed bioinformatics analysis, expressed and purified wild-type and slicing-defective AGO4 proteins from *Arabidopsis* and insect cells, and performed *in vitro* guide RNA loading and target RNA slicing assays. HYH and FW immunoprecipitated AGO4 in Figure 1E. HYH prepared small RNA sequencing libraries for Figure 2. JS performed RNA end-labeling for Figure 1F. JH prepared whole-genome bisulfite sequencing libraries and performed Illumina sequencing. FW and CSP wrote the manuscript.

## Supplemental Information

### Supplemental Materials and Methods

#### AGO4 immunoprecipitation from transgenic Arabidopsis plants

AGO4 proteins were immunoprecipitated from inflorescence tissues of transgenic *A. thaliana* plants, ecotype Wassilewskija (Ws), stably expressing FLAG-tagged wild-type AGO4 or slicing-defective AGO4-SD in the *ago4-4* mutant background. One gram of inflorescence tissue was ground to fine powder in liquid nitrogen and resuspended in 6 mL extraction buffer [50 mM Tris-HCl (pH 7.9), 75 mM sodium chloride, 5 mM magnesium chloride, 10% glycerol, 0.5% IGEPAL, 1 mM DTT, 1 mM PMSF and 1X plant protease inhibitor mix (Millipore Sigma)]. The homogenate was subjected to centrifugation at 16,000 g for 20 minutes at 4 °C. The supernatant was collected, filtered through a CellTrics 30 um strainer (Sysmex), and subjected to a second round of centrifugation at 16,000 g for 20 minutes at 4°C. The supernatant was incubated with 50 μL of anti-FLAG M2 agarose resin (Sigma) for 3 hours using a rotating mixer at 4 °C, followed by 4 washes, each with 1 mL of extraction buffer without plant protease inhibitors. The resin was resuspended in extraction buffer (without protease inhibitors) to a final volume of 100 μL. For protein analysis, 10 μL of the slurry containing immunoprecipitated AGO4 was boiled at 95 °C for 5 min in SDS sample buffer, resolved on 4-20% Mini-PROTEAN stain-free gels (Bio-Rad), and visualized by Coomassie Blue staining or immunoblotting with HRP-conjugated anti-FLAG M2 antibodies (Sigma) or anti-AGO4 polyclonal antibodies (Pikaard lab antibody stock #: 18187).

#### Extraction and analysis of AGO4-bound RNAs

Copurifying RNAs were released by incubating 50 μL of the slurry prepared as mentioned above at 70 °C for 5 min, precipitated with ethanol, and subjected to 5’ end-labeling with [γ-^32^P] ATP (Perkin Elmer) and T4 polynucleotide kinase (New England Biolabs). The labeled RNAs were passed through Performa Spin Columns (EdgeBio) to remove unincorporated radioactive ribonucleotides, resolved by polyacrylamide gel under denaturing condition (containing 7 M urea), and visualized by autoradiography. For small RNA sequencing, 25 μL of the slurry containing immunoprecipitated AGO4 was treated with 0.8 unit of proteinase K (New England BioLabs) followed by RNA extraction using TE saturated phenol/chloroform/isoamyl alcohol (pH 4.5). The RNA in the aqueous phase was ethanol precipitated, washed with 70% ethanol, and resuspended in nuclease-free water. The extracted RNAs were then used for high-throughput small RNA-seq library preparation.

#### Computational reconstitution of small RNA duplexes

Pairing patterns among siRNAs were analyzed using an in-house Python script. First, identical small RNAs aligning to the same genomic location were condensed into one sequence. Then, all possible pairs of 24 nt and 23 nt siRNAs that overlap in opposite orientation were analyzed. The distance from the 5’ end of a 5’A-bearing 24 nt siRNA to the 3’ end of the paired 23 nt siRNA (defined as 5’ to 3’ registry) for all 24/23 nt siRNA pairs was calculated and the frequencies of all possible pairing patterns were plotted. The 24/12 nt RNA pairing patterns were analyzed similarly. A simulated small RNA sequencing dataset mimicking the features of the actual small RNA-seq libraries was generated as a control using in-house script. To do so, the read alignment positions were randomized in the simulated libraries, while overall read abundance, read size distribution, and cluster locations were maintained. The simulated dataset was aligned to the *A. thaliana* TAIR10 genome assembly and analyzed in the same way as mentioned above. The in-house scripts for these analyses are available at: https://github.com/wangfeng3392/small_RNA.

#### *In vitro* slicing assay using immunoprecipitated AGO4 from Arabidopsis plants

AGO4 was immunoprecipitated from homozygous transgenic lines expressing either FLAG-AGO4 or slicing-defective FLAG-AGO4-SD in the *ago4-4* null mutant background. To prepare AGO4 immunoprecipitation fractions for 8 slicing reactions, two grams of inflorescence tissue was ground to a fine powder in liquid nitrogen and resuspended in 6 mL extraction buffer [400 mM Potassium acetate, 25 mM HEPES-KOH (pH 7.4), 5 mM Magnesium acetate, 10% glycerol, 0.1% IGEPAL, 0.5 mM DTT, 1 mM PMSF and 1X plant protease inhibitor mix (Sigma)]. The homogenate was subjected to centrifugation at 16,000 g for 20 minutes at 4 °C. The supernatant was collected, filtered through CellTrics 30 um (Sysmex), and subjected to a second round of centrifugation at 16,000 g for 20 minutes at 4°C.

FLAG antibody-conjugated Dynabeads for 8 reactions were prepared with the following method: 64 µL Protein G-conjugated Dynabeads (Thermo Fisher Scientific) were incubated with 16 µL monoclonal anti-FLAG M2 antibody (F1804, Millipore Sigma) for no less than 30 minutes at room temperature, or overnight at 4 °C. Excess antibody was removed by pulling down Dynabeads using a DynaMag magnet (Thermo Fisher Scientific). The supernatant was discarded, and the beads were washed 3 times with 1 mL extraction buffer [400 mM Potassium acetate, 25 mM HEPES-KOH (pH 7.4), 5 mM Magnesium acetate, 10% glycerol, 0.1% IGEPAL, 0.5 mM DTT, 1 mM PMSF and 1X plant protease inhibitor mix (Sigma)]. To precipitate AGO4 proteins, antibody-conjugated beads were incubated with clarified lysate for 2 hours at 4 °C with constant rotation. After removing the lysate, the beads were washed three times with wash buffer [600 mM Potassium acetate, 25 mM HEPES-KOH (pH 7.4), 5 mM Magnesium acetate, 10% glycerol, 0.1% IGEPAL, 1 mM PMSF], and then washed once with reaction buffer [100 mM Potassium acetate, 25 mM HEPES-KOH (pH 7.4), 5 mM Magnesium acetate, 10% glycerol, 0.01% IGEPAL, and 1 mM PMSF]. For each wash, the beads were resuspended by pipetting up and down followed by rotating for 5 minutes at 4 °C. The beads were then pulled down using the DynaMag magnet, resuspended in 160 µL reaction buffer containing 1 unit/μL RiboLock RNase Inhibitor (Thermo Fisher), and evenly aliquoted to 8 tubes.

Non-radioactive phosphorylated guide RNA was added to AGO4 loading reactions to a final concentration at 40 nM. The reactions were incubated at room temperature for 1 hour with constant rotation. The beads were then magnet-precipitated and washed three times with reaction buffer to remove unbound guide RNAs. The beads were then resuspended in 20 μL reaction buffer containing 1 unit/μL RiboLock RNase Inhibitor (Thermo Fisher), ^32^P-phosphorylated target RNA (∼1000 CPM per reaction), and 1 µg yeast tRNAs (Thermo Fisher Scientific). The reactions were further incubated for 1 hour at room temperature with constant rotation. After incubation, Dynabeads were magnet-precipitated. The supernatant fraction was transferred to a fresh tube. The beads were subject to 3 washes with reaction buffer followed by magnet-precipitation (the bead fraction). The reactions in both supernatant and bead fractions were quenched by adding 200 µL of stop solution (300 mM sodium acetate and 7M urea). RNAs were extracted with 200 µL phenol/chloroform/isoamyl alcohol followed by ethanol precipitation. The resulted RNA pellets were washed once with 70% ethanol, allowed to dry, and resuspended with 1X formamide loading buffer [40% deionized formamide, 0.5 mg/mL xylene cyanol, 0.5 mg/ml bromophenol blue, and 5 mM EDTA (pH 8.0)]. The resuspended RNA samples were then denatured at 95 °C and resolved in a 15% polyacrylamide gel containing 7M urea by electrophoresis with a constant wattage of 50 W. The gels were dried and visualized by phosphorimaging.

#### Recombinant AGO4 expression and purification

Codon-optimized N-terminal FLAG-tagged AGO4 or slicing-defective AGO4-SD (with mutations D660A, D742A, and H874A) were first sub-cloned into SUMOstar Insect Intracellular Vector (LifeSensors), and then transformed into *E. coli* strain DH10Bac to generate bacmid DNA. Baculovirus was prepared as previously described (Fitzgerald et al., 2006) except that Sf9 cells cultured in Sf-900 II serum-free media (Thermo Fisher Scientific) was used for virus production. To express recombinant AGO4 and slicing-defective AGO4-SD proteins, 2 L of Sf9 cells with a cell density of approximately 1.5X 10^6^ cells/mL were infected at a multiplicity of infection (MOI) of 2 and further cultured on a shaker at 125 rpm for 48 to 60 hours, at 27 °C. Cells were collected by centrifuge at 800 g for 10 minutes, washed once with 1X phosphate buffered saline (PBS) buffer, pelleted again at 800 g for 10 minutes, and flash frozen in liquid nitrogen. All subsequent purification steps were carried out at 4 °C. Cell pellets were resuspended in 400 mL ice-cold lysis buffer [400 mM Potassium acetate, 25 mM HEPES-KOH (pH 7.4), 5 mM Magnesium acetate, 10% glycerol, 0.1% IGEPAL, 0.5 mM DTT, 1 mM PMSF and 1X plant protease inhibitor mix (Sigma)]. Cells were disrupted by 8 strokes in a prechilled Dounce Homogenizer. The lysate was clarified by centrifugation at 48,000 g for 45 min followed by immunoprecipitation using 2 mL pre-equilibrated anti-FLAG M2 agarose beads (Sigma) for 2 hours. The beads were subject to 3 washes, each time using 10 mL of wash buffer [400 mM Potassium acetate, 25 mM HEPES-KOH (pH 7.4), 5 mM Magnesium acetate, 10% glycerol, 0.1% IGEPAL, and 1 mM PMSF]. Recombinant AGO4 was eluted 3 times by incubating the beads with 2 mL wash buffer containing 0.5 mg/mL 3X FLAG peptide (APEXBIO) and 20 mM Imidazole. The eluted fraction was further incubated with 0.5 mL of Nickel-NTA agarose beads (Qiagen) for 1 hour. The beads were washed 3 times in the wash buffer containing 40 mM Imidazole. Recombinant AGO4 was eluted 3 times using NiNTA elution buffer [250 mM Imidazole, 400 mM Potassium acetate, 25 mM HEPES-KOH (pH 7.4), 5 mM Magnesium acetate, 10% glycerol, 0.01% IGEPAL, and 1 mM PMSF]. The pooled eluted fraction was dialyzed into storage buffer [150 mM Potassium acetate, 25 mM HEPES-KOH (pH 7.4), 5 mM Magnesium acetate, 10% glycerol, 0.01% IGEPAL, 0.5 mM DTT, 1 mM PMSF], concentrated to ∼ 500 ng/µL, divided to 8 µL aliquots, and flash frozen in liquid nitrogen.

#### *In vitro* slicing using insect cell expressed recombinant AGO4

Recombinant AGO4 or AGO4-SD (approximately 1 µg each) were incubated with 40 nM of unlabeled phosphorylated guide RNA in 20 µL of reaction buffer containing 1 unit/μL RiboLock RNase Inhibitor (Thermo Fisher) for 1 hour at room temperature to allow guide RNA incorporation into AGO4. 1 µL ^32^P-phosphorylated target RNA (∼1000 cpm/µL) and 1 µg yeast tRNA (Thermo Fisher Scientific) was added to each reaction. After 1 hour incubation at room temperature, reactions were quenched by adding 200 µL of stop solution (300 mM NaOAc and 7M urea). RNAs were extracted by 200 µL of phenol/chloroform/isoamyl alcohol, ethanol precipitated, resuspended in 1X formamide loading buffer, resolved in 15% polyacrylamide gels containing 7M urea at a constant wattage of 50 W, and visualized by phosphorimaging.

For the test of guide RNA paired with a 12 nt passenger strand fragment, 1 µM of phosphorylated guide RNA was mixed with 2 µM of phosphorylated 12 nt passenger fragment in 50 µL of annealing buffer [100 mM Potassium acetate, 25 mM HEPES-KOH (pH 7.4), 5 mM Magnesium acetate, and 1 unit/μL RiboLock RNase Inhibitor (Thermo Fisher)] in a microcentrifuge tube. Annealing was achieved by heating the oligonucleotide mixture in a boiling water bath and then allowing the water bath to cool to room temperature. Recombinant AGO4 (approximately 1 µg) was incubated with annealed RNAs containing 40 nM guide-strand and 80 nM 12 nt fragment in 20 µL of reaction buffer containing 1 unit/μL RiboLock RNase Inhibitor (Thermo Fisher) for 1 hour at room temperature to allow RNA incorporation into AGO4. 1 µL ^32^P-phosphorylated target RNA (∼1000 cpm/µL) and 1 µg yeast tRNA (Thermo Fisher Scientific) was then added to each reaction. After 1-hour incubation at room temperature, reactions were quenched by adding 200 µL of stop solution (300 mM NaOAc and 7M urea). RNAs were extracted using 200 µL of phenol/chloroform/isoamyl alcohol, ethanol precipitated, resuspended in 1X formamide loading buffer, resolved in 15% polyacrylamide gels containing 7M urea at a constant wattage of 50 W, and visualized by phosphorimaging.

#### *In vitro* guide RNA loading assay

To test if guide siRNAs with 5’ A, U, G, and C can be incorporated into AGO4 (Figure 4D), recombinant AGO4 (approximately 1 µg) was incubated with 10 nM ^32^P-phosphorylated guide RNAs in 20 µL of reaction buffer containing 1 unit/μL RiboLock RNase Inhibitor (Thermo Fisher) for 1 hour at room temperature to allow guide RNA loading. AGO4 proteins were then subjected to immunoprecipitation using 8 µL of anti-FLAG antibody-conjugated Dynabeads. Following 1 hour of incubation at room temperature, the beads were magnet-precipitated and washed three times with 0.5 mL wash buffer [600 mM Potassium acetate, 25 mM HEPES-KOH (pH 7.4), 5 mM Magnesium acetate, 10% glycerol, 0.1% IGEPAL, 0.5 mM DTT, and 1 mM PMSF]. Reactions were quenched by adding 200 µL of stop solution (300 mM NaOAc and 7M urea). RNA was extracted by 200 µL phenol/chloroform/isoamyl alcohol, ethanol precipitated, resuspended in 1x formamide loading buffer, and subject to electrophoresis in 15% polyacrylamide gels containing 7M urea at a constant wattage of 50 W. The gels were dried and visualized by phosphorimaging. For competition assays, cold phosphorylated guide RNA was mixed with ^32^P-phosphorylated guide RNA that contains a 5’ adenosine. The mixed RNA was then subjected to AGO4 loading, immunoprecipitation, and electrophoresis as described above.

#### RNA blot analysis

Recombinant AGO4 or AGO4-SD (approximately 1 µg each) was incubated with 40 nM of guide siRNA, having either 5’ monophosphate, 5’ hydroxyl, or 5’ triphosphate groups, in 20 µL of reaction buffer containing 1 unit/μL RiboLock RNase Inhibitor (Thermo Fisher) for 1 hour at room temperature. AGO4-siRNA complexes were immunoprecipitated by incubating with 8 µL of FLAG antibody-conjugated Dynabeads for 2 hours at 4 °C. The beads were then washed twice with 0.5 mL of wash buffer. Reactions were quenched by adding 200 µL of stop solution containing 300 mM NaOAc and 7M Urea to the beads. RNAs were extracted by 200 µL phenol/chloroform/isoamyl alcohol, ethanol precipitated, resuspended in 1X formamide loading buffer, and resolved in a 15% polyacrylamide gel containing 7M urea. RNAs were then transferred to a nylon membrane (Amersham Hybond N+) using 0.5X TBE buffer in a semi-dry blotter (Bio-Rad) at constant voltage of 20 V for 45 minutes. The RNA was then chemically crosslinked to the membrane with 31 mg/mL of 1-ethyl-3-(3-dimethylaminopropyl)-carbodiimide (EDC) for 2 hours at 60 °C. The membrane was washed 5 times with water, followed by prehybridization using PerfectHyb Plus buffer (Sigma) containing 100 µg /mL sheared salmon sperm DNA (Ambion) at 42 °C for 1 hour. The membrane was then hybridized to 1 µM 5’-end ^32^P-labeled DNA probe overnight in PerfectHyb Plus buffer containing 100 µg/mL sheared salmon sperm DNA at 42 °C. Following hybridization, the membrane was washed twice with non-stringent RNA blot wash buffer [3x SSC, 25 mM NaH2PO4 (pH 7.5), and 5% SDS] at 50 °C for 10 minutes, twice with non-stringent RNA blot wash buffer at 50 °C for 30 minutes, followed by one wash with stringent RNA blot wash buffer (1X SSC and 1% SDS) at 50 °C for 5 minutes. The membrane was analyzed by phosphorimaging.

## Supplemental Figure Legends

**Figure S1.**
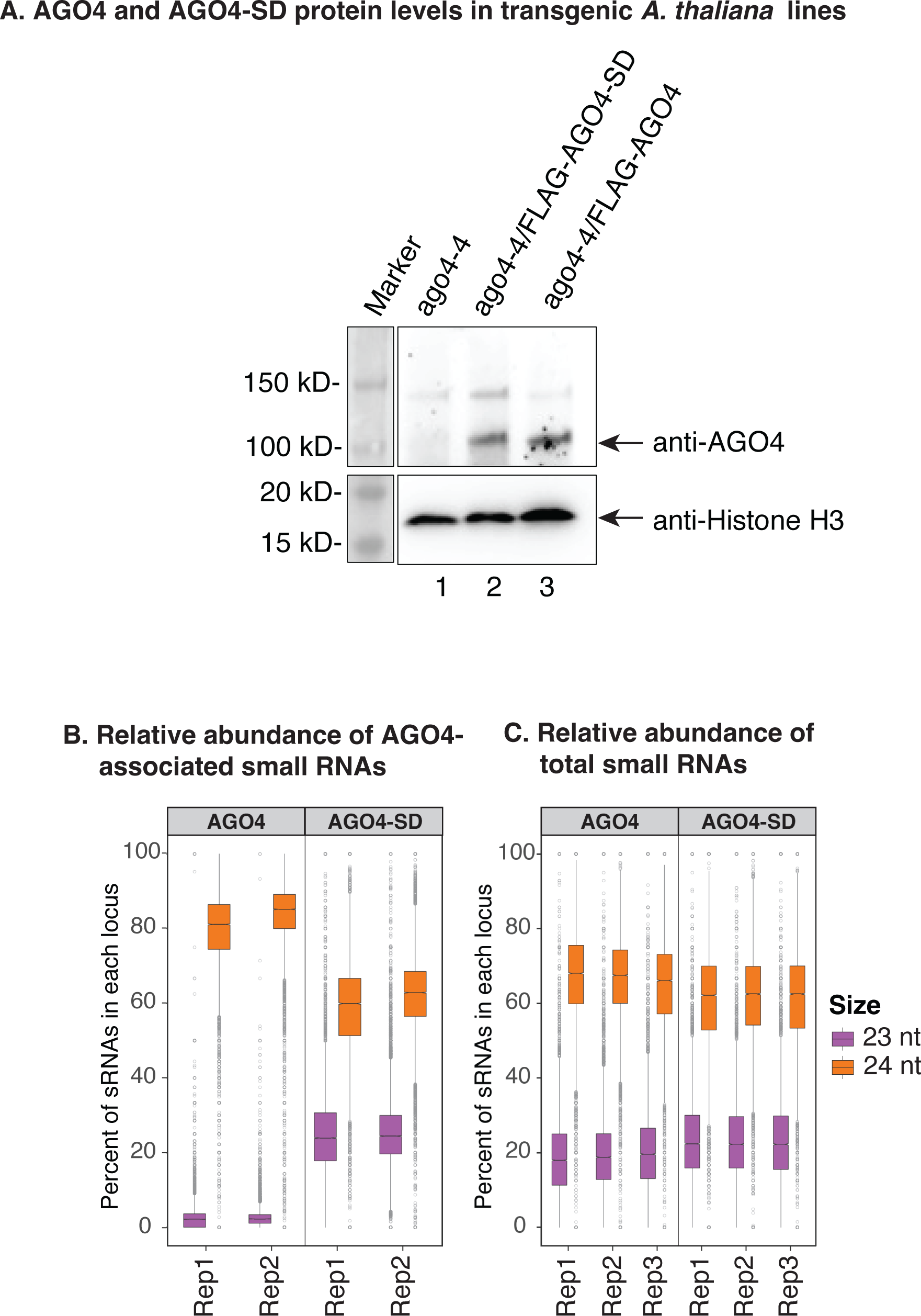
Relative abundance of 23 and 24 nt RNAs associated with AGO4 or in the total small RNA pool. Related to Figures 2A and 2B. **A.** AGO4 and AGO4-SD protein levels in transgenic *A. thaliana* lines. For each line,100 mg of 10-day-old seedling tissue was ground into fine powder in liquid nitrogen then resuspended in 200 µL of 2X SDS-PAGE sample buffer. 30 µL of each sample was then loaded onto a 4-20% SDS-PAGE gel, resolved by electrophoresis and subjected to immunoblotting. Anti-FLAG antibody (A8592) was used to detect FLAG-tagged AGO4 or AGO4-SD proteins. Histone H3 proteins were detected using anti-Histone H3 antibody (ab21054). **B.** Relative abundance of 23 nt and 24 nt siRNAs co-immunoprecipitated with wild-type AGO4 or slicing-defective AGO4-SD at loci at which 24 nt siRNAs predominate. Boxplots show medians (horizontal lines), 1st–3rd quartile range (boxes), other data extending to 1.5 times the interquartile range (whiskers), and outliers for two independent replicates. **C**. Relative abundance of 23 nt and 24 nt siRNAs in the pool of total RNAs purified from inflorescence tissues. Total RNAs were extracted from stable transgenic lines expressing wild-type or slicing-defective AGO4 in the *ago4-4* null mutant background. Results of three independent replicates are shown.

**Figure S2.**
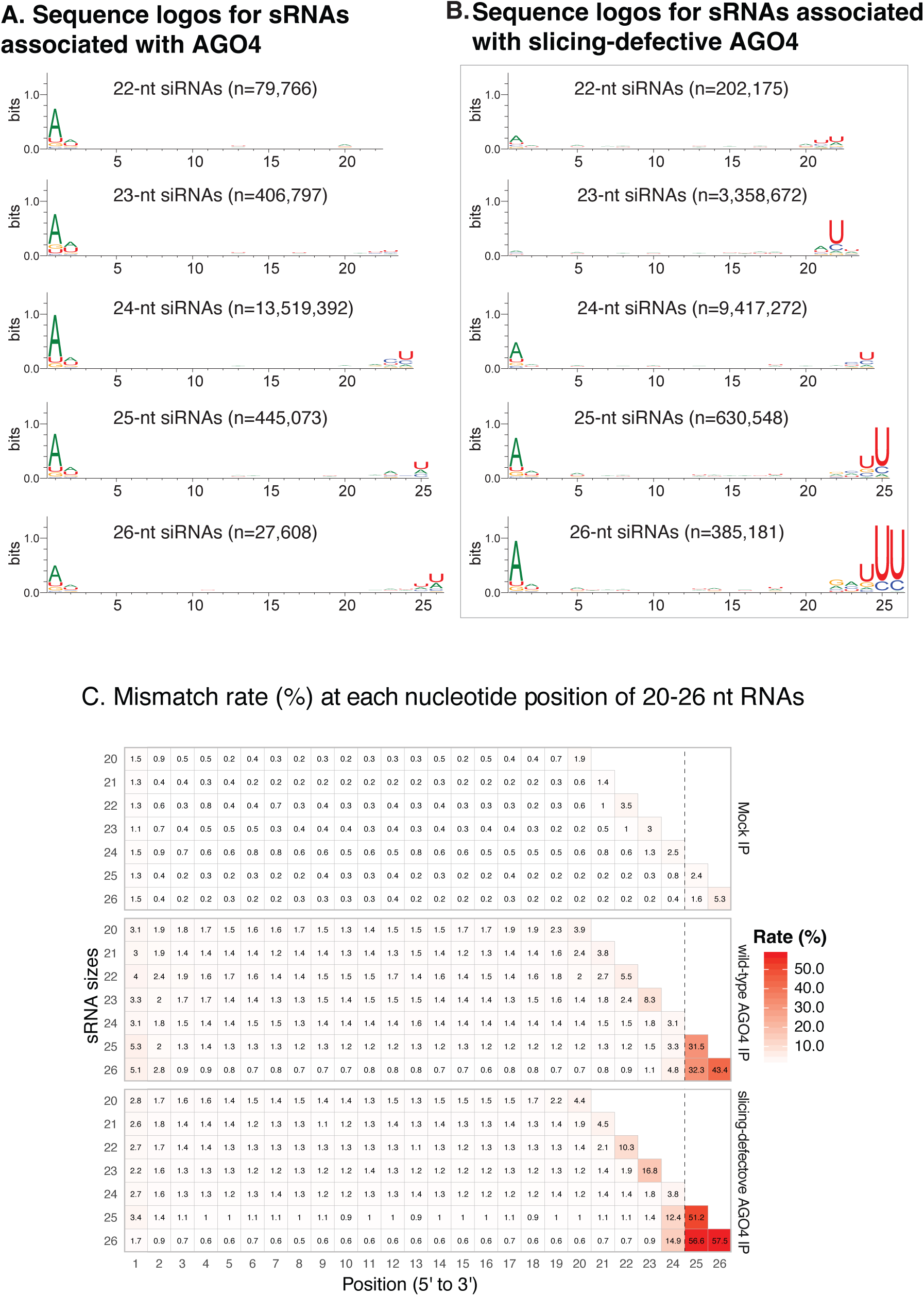
Sequence features of AGO4-associated small RNAs. Related to Figures 2A and 2B. **A**. Sequence logos for 22-26 nt small RNAs that co-immunoprecipitated with wild-type AGO4. **B**. Sequence logos for 22-26 nt small RNAs that co-immunoprecipitated with slicing-defective AGO4-SD. **C**. Genome mismatch frequency for each nucleotide position of 22-26 nt small RNAs that co-immunoprecipitated (IPed) with wild-type AGO4 or slicing-defective AGO4-SD or the mock IP control.

**Figure S3.**
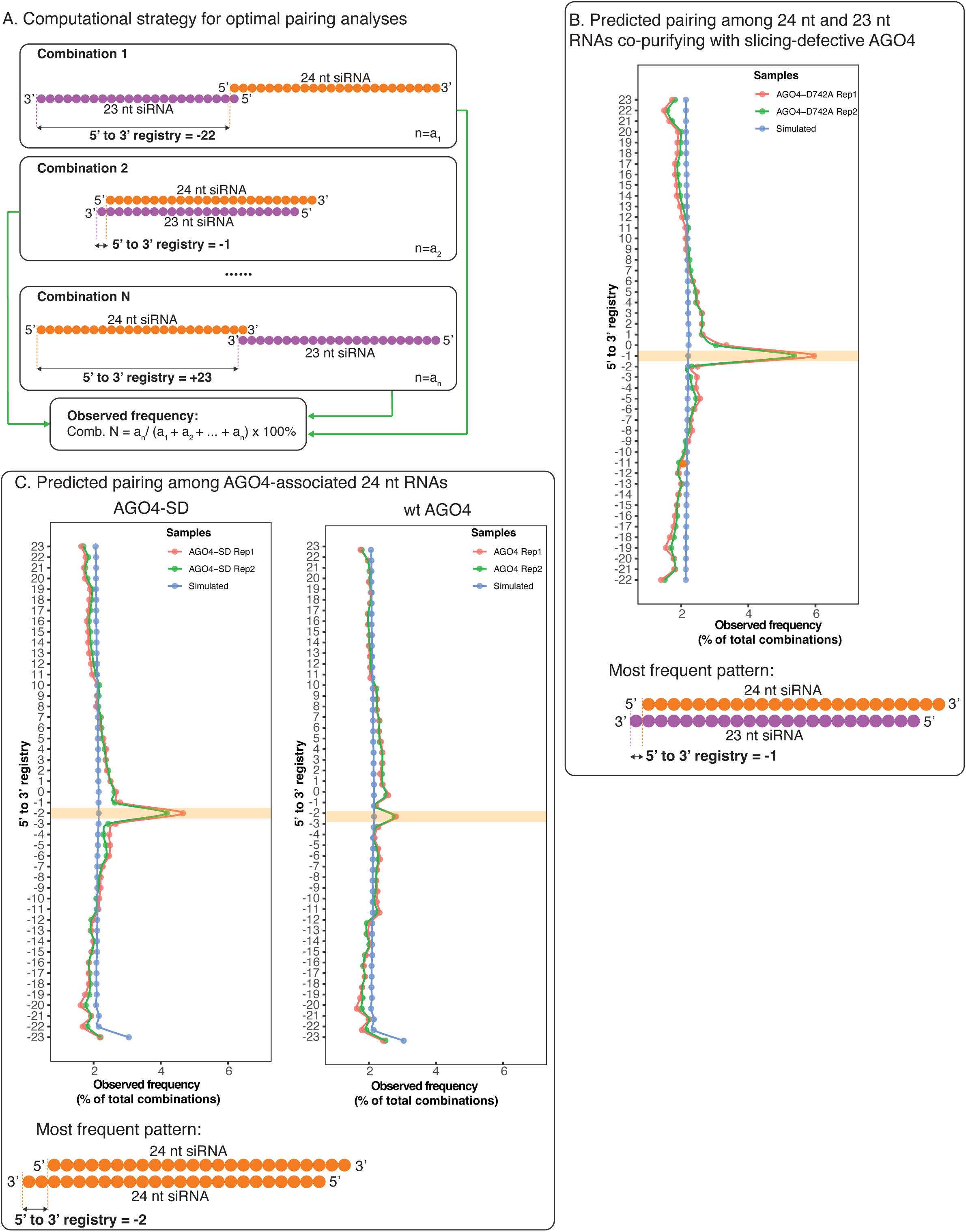
Computational reconstruction of 24/23 or 24/24 siRNA duplexes associated with slicing-defective AGO4. Related to Figures 2C and 2D. **A**. Computational strategy for prediction of AGO4-bound siRNA duplexes. RNAs associated with slicing-defective AGO4, thus representing both guide and passenger strands, were tested for optimal basepairing alignments. For 24 nt and 23 nt siRNAs that overlap with at least 1 nucleotide of complementarity, the 5’ to 3’ registry is defined as the distance from the 5’ nucleotide of the top strand to the 3’ nucleotide of the bottom strand. The frequency of every possible 5’ to 3’ registry in the population of all observed 24/23 nt pairs was calculated. **B**. Predicted pairing frequencies represented by the possible 5’ to 3’ registries for 24/23 nt siRNA pairs. The most highly represented pairing pattern is illustrated at the bottom of the panel. **C**. Predicted pairing frequencies represented by the possible 5’ to 3’ registries for 24/24 nt siRNA pairs in AGO4-SD and wild-type AGO4. The most highly represented pairing pattern for 24 nt RNAs copurifying with slicing-defective AGO4 is illustrated at the bottom of the panel.

**Figure S4.**
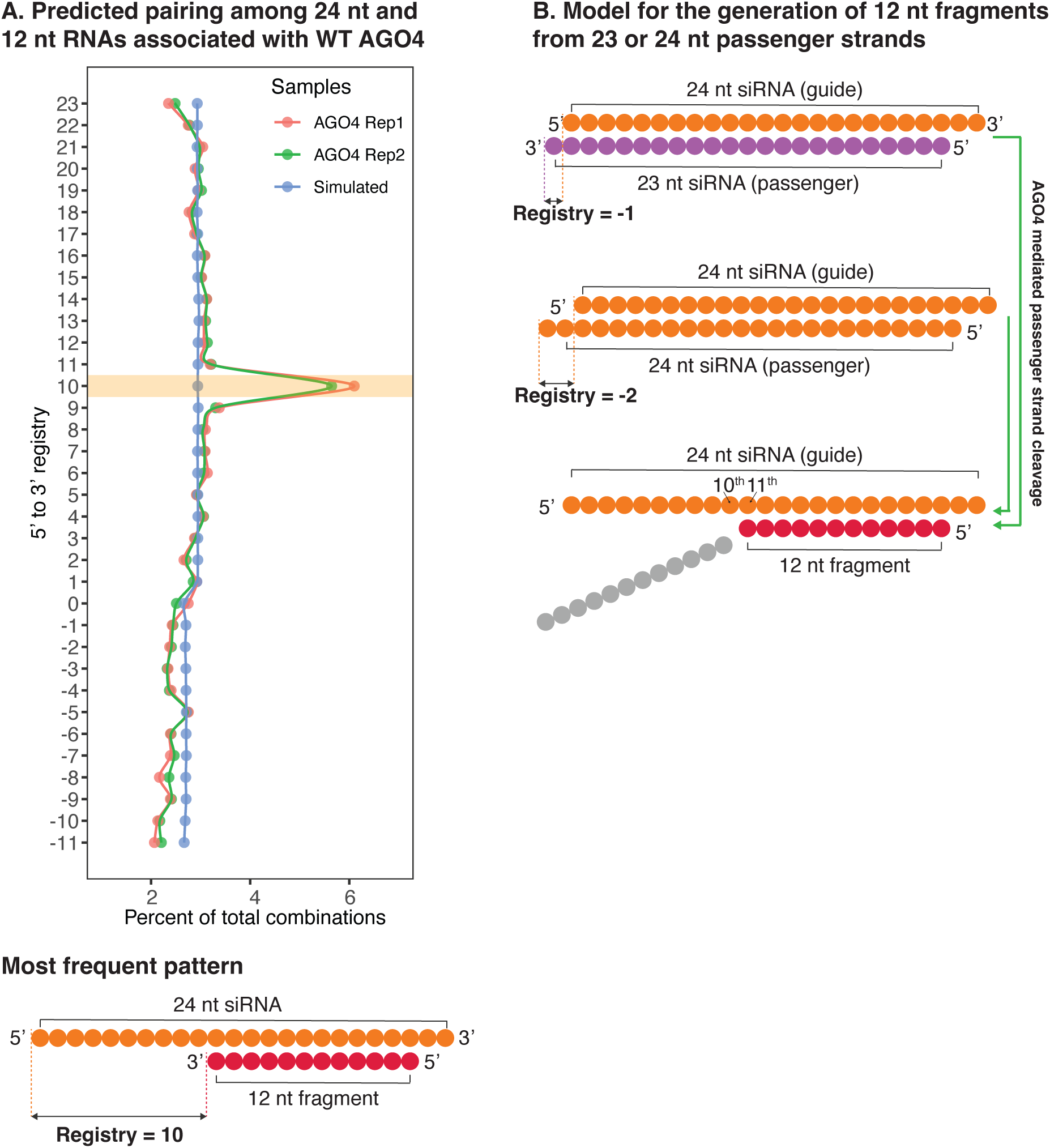
Computational prediction of pairing between 12 nt and 24 nt RNAs associated with wild-type AGO4. Related to Figures 2C and 2D. A. Using the same computational approach used in Figure S3, the plot of the frequencies of various pairing registries for 12 and 24 nt RNAs associated with wild-type AGO4 is shown, with the most frequently represented pairing pattern displayed at the bottom of the panel. B. Model for how slicing generates 12 nt fragments from passenger strands that are either 23 or 24 nt in length. In both cases, AGO4 slicing of the passenger strand at a position complementary to the 10th nucleotides of the 24 nt guide strand, counting from the 5’ end, accounts for 12 nt RNAs.

**Figure S5.**
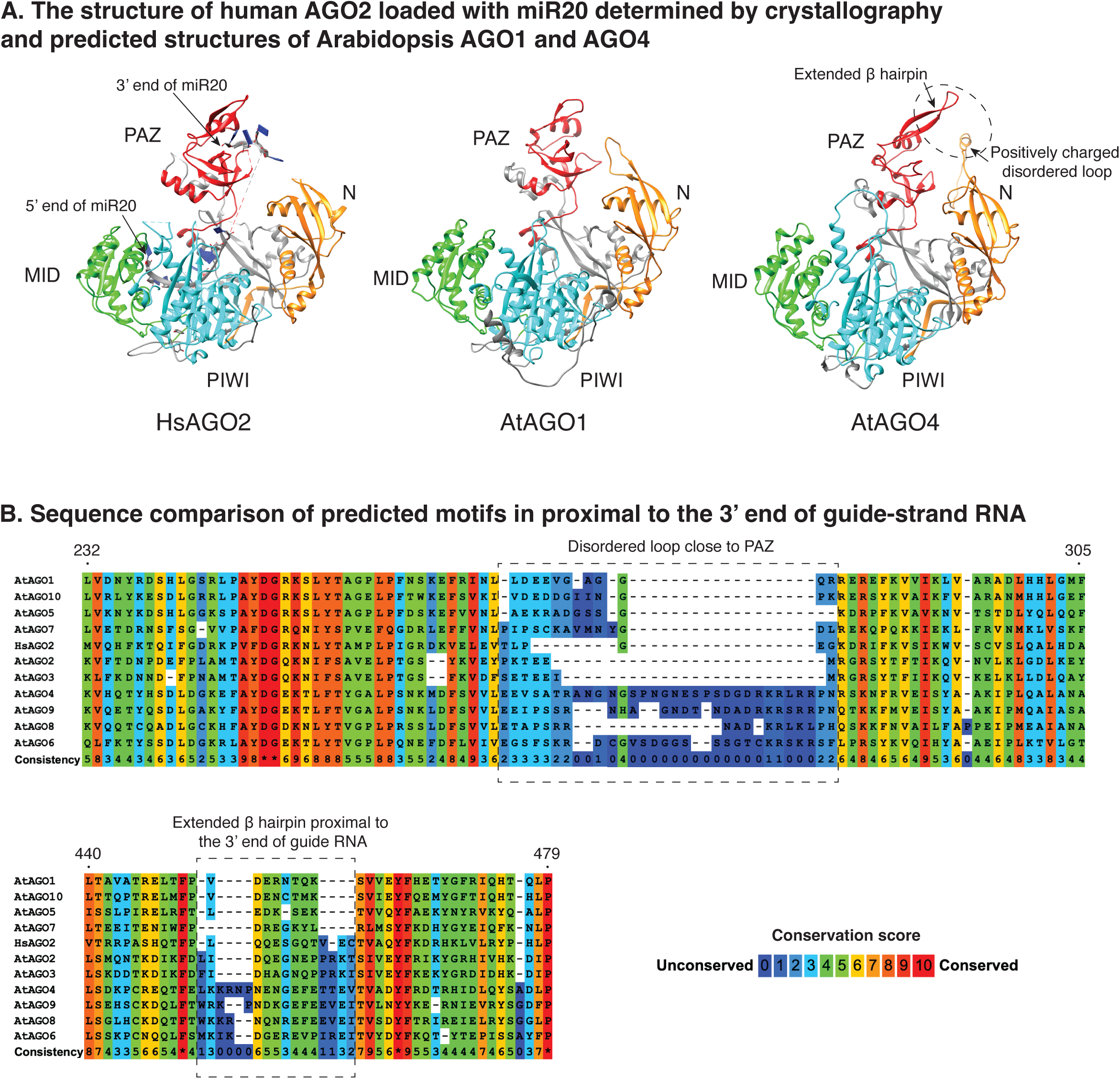
Comparison of the human AGO2 structure to the predicted structures for *A. thaliana* AGO1 and AGO4 in the region near the guide strand RNA’s 3’ end. Related to Figure 3C. A. Crystal structure for human AGO2 in association with miR20 (PDB: 4F3T) and AlphaFold-predicted structures for *A. thaliana* AGO1 and AGO4. An extended b hairpin and positively charged disordered loop in the predicted structure of AGO4 is highlighted. B. Sequence comparisons of predicted motifs predicted to be in proximity to guide strand RNA 3’ ends are shown for the ten A*. thaliana* AGO proteins and human AGO2. Residues are color-coded based on degree of conservation, with warmer colors representing higher conservation.

**Figure S6.**
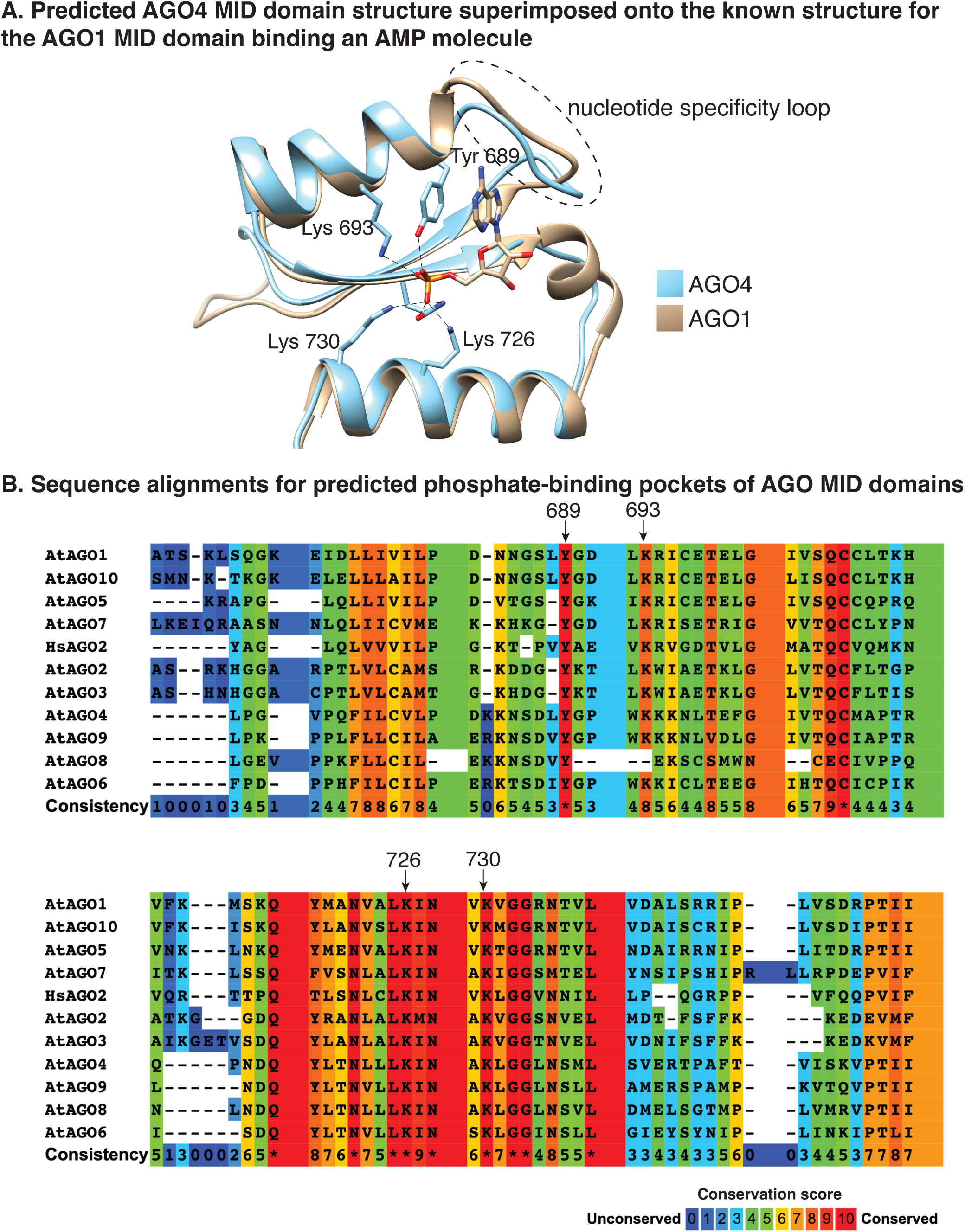
Comparison of the AGO1 phosphate binding pocket for the guide strand’s 5’ end to the equivalent structure predicted for AGO4. Related to Figure 4B and 4C. A. The MID domain of *Arabidopsis* AGO4 predicted using AlphaFold is superimposed onto the crystal structure for the *Arabidopsis* AGO1 MID domain in complex with the phosphate group of Adenosine monophosphate (PDB: 4G0Y). Residues in AGO1 predicted to contact the 5’ phosphate are highlighted. Predicted hydrogen bonds are denoted by dashed lines. The nucleotide specificity loop is denoted by a dashed circle. The numbering of highlighted amino acids is based on the *Arabidopsis* AGO1 sequence. B. Sequence comparison of the predicted phosphate-binding pocket in the ten *A. thaliana* AGO proteins and human AGO2. Key residues predicted to contact the 5’ terminal phosphate are highlighted. Residues are color-coded based on conservation, with warmer color represents more highly conserved sequences.

**Figure S7.**
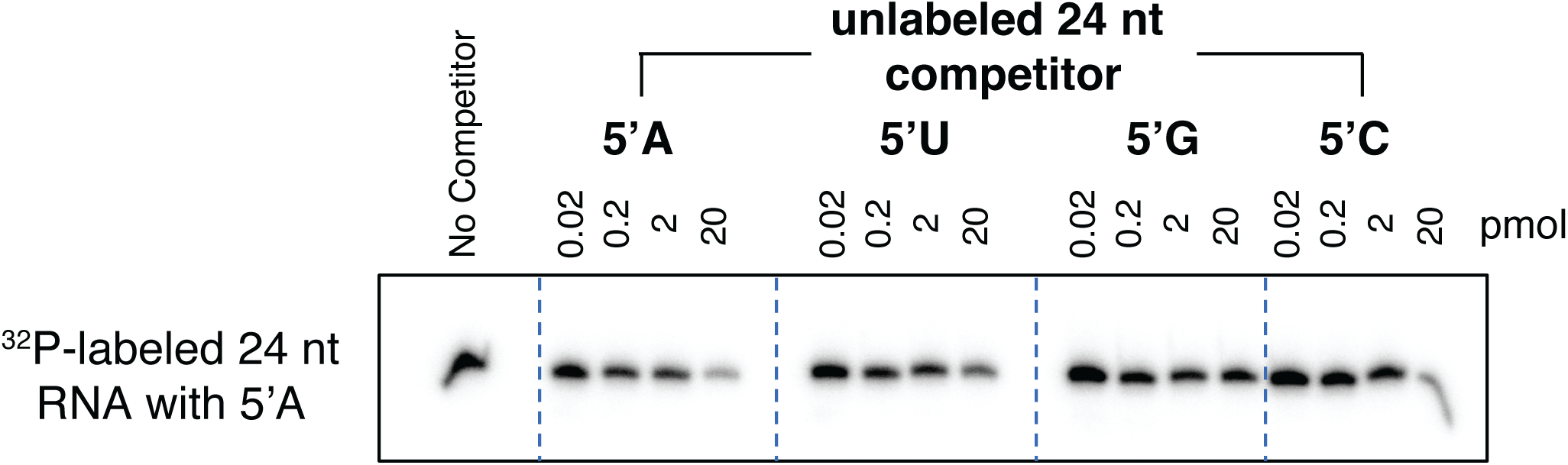
Competition assays testing whether the 5’ terminal nucleotide of 24 nt guide RNAs affects AGO4 binding. Related to Figures 4D and 4E. FLAG-tagged AGO4 binding to a ^32^P end-labeled 24 nt siRNA beginning with a 5’ A was carried out in the presence of increasing concentrations of unlabeled phosphorylated 24 nt siRNAs that begin with 5’ A, U, G, or C but are otherwise identical in sequence. AGO4 was then affinity captured and associated RNAs were resolved by denaturing PAGE and autoradiography.

**Figure S8.**
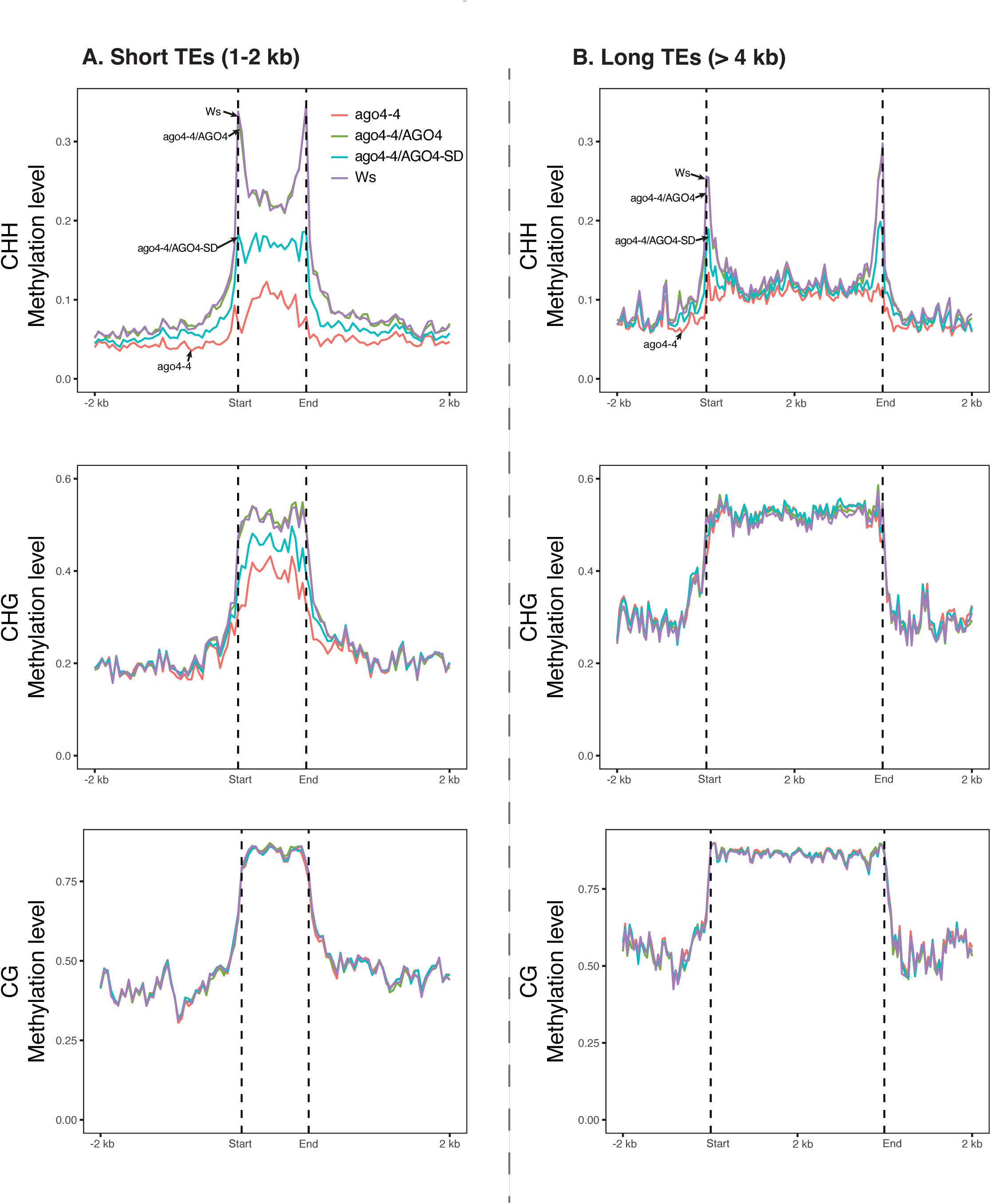
Influence of AGO4 slicing activity on CHH, CHG, and CG methylation at transposable elements. Arabidopsis short TEs (1-2 kb) and long TEs (>4kb) that overlap with AGO4-dependent DMRs were aligned at their 5’ and 3’ ends. Average cytosine methylation for all cytosines within 50 bp intervals in TE bodies and 2 kb upstream and downstream flanking regions is plotted.

Table S1. Genomic positions for AGO4-associated RNA clusters

Table S2. CHH methylation ratios at AGO4 Differentially Methylated Regions (DMRs)

**Table S3.**
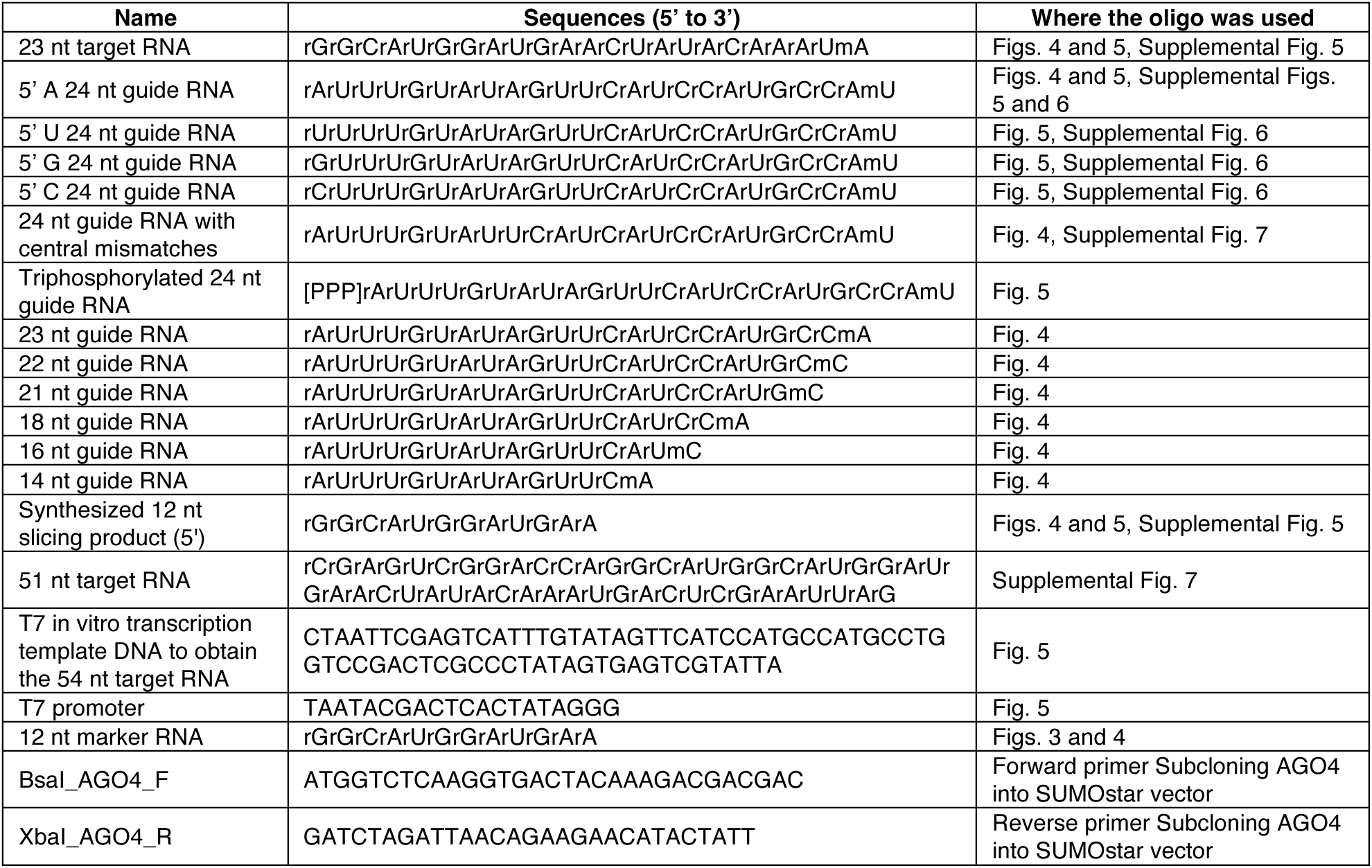
Oligonucleotides used in the study.

